# Characterization of the REC114-MEI4-IHO1 complex regulating meiotic DNA double-strand break formation

**DOI:** 10.1101/2023.01.11.523614

**Authors:** Hamida Laroussi, Ariadna B. Juarez-Martinez, Aline Le Roy, Elisabetta Boeri Erba, Bernard de Massy, Jan Kadlec

## Abstract

Meiotic recombination is initiated by the formation of DNA double-strand breaks (DSBs), essential for fertility and genetic diversity. In the mouse, DSBs are formed by the catalytic TOPOVIL complex consisting of SPO11 and TOPOVIBL. To preserve genome integrity, the activity of the TOPOVIL complex is finely controlled by several meiotic factors including REC114, MEI4 and IHO1, but the underlying mechanism is poorly understood. Here, we report that mouse REC114 forms homodimers, that it associates with MEI4 as a 2:1 heterotrimer that further dimerizes, and that IHO1 forms coiled-coil based tetramers. Using AlphaFold2 modelling combined with biochemical characterization we uncovered the molecular details of these assemblies. Finally, we show that IHO1 directly interacts with the PH domain of REC114 by recognizing the same surface as TOPOVIBL and another meiotic factor ANKRD31. These results provide strong evidence for the existence of a ternary IHO1-REC114-MEI4 complex and show that REC114 is a potential regulatory platform mediating mutually exclusive interactions with several partners.

## Introduction

At the onset of prophase of the first meiotic division, a unique program of DNA double-strand break (DSB) formation takes place (de Massy, 2013; Lam and Keeney, 2015). Ten to a hundred DSBs in each meiotic cell are formed at preferred DNA sites, named hotspots (Tock and Henderson, 2018). The DSBs are repaired by homologous recombination generating connections between paternal and maternal chromosomes that are essential for proper chromosome segregation at the first meiotic division (Baudat et al., 2013; Hunter, 2015). The meiotic DSB formation is evolutionary conserved and has to be carefully controlled to preserve the genome integrity.

The key player in the meiotic DSB formation is the catalytic complex TOPOVIL, in the mouse consisting of SPO11 and TOPOVIBL (Baudat et al., 2000; Robert et al., 2016; Romanienko and Camerini-Otero, 2000) The TOPOVIL complex is overall related to the archaeal TopoVI type IIB topoisomerases (Brinkmeier et al., 2022) but the mouse TOPOVIBL subunit differs from TopoVIB, in particular it lacks its ATP-biding and dimerization sites (Nore et al., 2022). The activity of the catalytic complex is regulated by several meiotic factors. In the mouse, these include REC114, MEI4, IHO1 and MEI1 that are essential for DSB formation and localize as foci on chromosome axes at meiotic prophase onset (Acquaviva et al., 2020; Dereli et al., 2021; Kumar et al., 2018, 2015, 2010; Reinholdt and Schimenti, 2005; Stanzione et al., 2016).

REC114 consists of 259 amino acid residues. In its N-terminus it possesses a Pleckstrin homology (PH) domain whose crystal structure was determined (Boekhout et al., 2019; Kumar et al., 2018; Nore et al., 2022). The C-terminal part of REC114 directly interacts with the N-terminus of MEI4 (Kumar et al., 2018). REC114 also interacts in yeast two-hybrid assay (Y2H) with IHO1. These interactions may predict a tripartite REC114-MEI4-IHO1 complex but it has not been identified yet. Since IHO1 also binds the unsynapsed chromosomal axis protein HORMAD1 (Stanzione et al., 2016), it has been hypothesized that a putative REC114-MEI4-IHO1 complex called pre-DSB recombinosome might assemble along the HORMAD1-containing chromosomal axis and regulate the TOPOVIL complex activity (Stanzione et al., 2016). Along these lines, the PH domain of REC114 was recently shown to directly interact with the C-terminal peptide of TOPOVIBL (Nore et al., 2022). Structure-based mutations that disrupt this interaction strongly reduce the DSB activity genome-wide in oocytes, while only in sub-telomeric regions in spermatocytes (Nore et al., 2022). The PH domain of REC114 also interacts with a C-terminal fragment of another meiotic factor ANKRD31 (Boekhout et al., 2019) which is involved in regulating DSB number and localization, and this interaction is mutually exclusive with that of TOPOVIBL (Nore et al., 2022; Papanikos et al., 2019). The exact role and the interplay of these factors in DSB formation still remain unclear.

In *S. cerevisiae* the proteins and mechanism involved in DSB formation have been analysed at the genetic and biochemical level (Yadav and Claeys Bouuaert, 2021). The TOPOVIBL subunit of the catalytic complex exists in yeast as Rec102/Rec104 dimer (Arora et al., 2004; Claeys Bouuaert et al., 2021b; Jiao et al., 2003). The *S. cerevisiae* core complex consists of Spo11, Rec102, Rec104 and an additional protein Ski8 (Claeys Bouuaert et al., 2021b). The accessory proteins Rec114–Mei4 and Mer2 (IHO1 ortholog) form two separate complexes in vitro with 2:1 and tetrameric stoichiometry, respectively (Claeys Bouuaert et al., 2021a; Rousova et al., 2021). The two complexes bind DNA with high cooperativity and assemble into condensates that might recruit the catalytic complex to DNA through an Y2H-observed interaction of Rec114 with Rec102/Rec104 (Arora et al., 2004; Claeys Bouuaert et al., 2021a; Maleki et al., 2007). Since the PH domain surface interacting with TOPOVIBL in mouse is not conserved in yeast, the molecular details of this interaction might differ from the mouse REC114-TOPOVIBL interaction (Nore et al., 2022). While counterparts of most mammalian meiotic factors exist in yeast, their sequence conservation is often very low, indicating possible differences in the molecular details underlying their activities.

Here, using biochemical and biophysical analyses combined with AlphaFold2 modelling, we uncovered molecular details of the interaction between mouse REC114 and MEI4, where two molecules of REC114 interact with their C-termini with the N-terminal helix of MEI4 forming 2:1 and 4:2 assemblies. We also show that an equivalent REC114 homodimeric complex is formed even in absence of MEI4. In addition, we found IHO1 that forms coiled-coil based tetramers and interacts specifically via its N-terminus with the PH domain of REC114, in a way that is incompatible with REC114 binding to ANKRD31 and TOPOVIBL. Together, these results provide strong evidence for the existence of a complex between mouse IHO1 and REC114-MEI4, with a possible stoichiometry of 4IHO1:4REC114:2MEI4.

## Results

### Characterization of the REC114-MEI4 complex

REC114 interacts directly with MEI4 in pull-down experiments and residues 203-254 of REC114 and 1-127 of MEI4 are sufficient for the interaction (Kumar et al., 2018) (Fig. 1a). Using Strep-tag pull-down assays, with co-expressed proteins, we could show that Strep-REC114 further truncated to residues 226-254 can still bind MEI4 (Fig. 1b). Similarly, MEI4 encompassing only the N-terminal 43 residues was sufficient for the interaction with co-expressed His-MBP-REC114 in an MBP pull-down assay (Fig. 1c).

**Figure 1.**
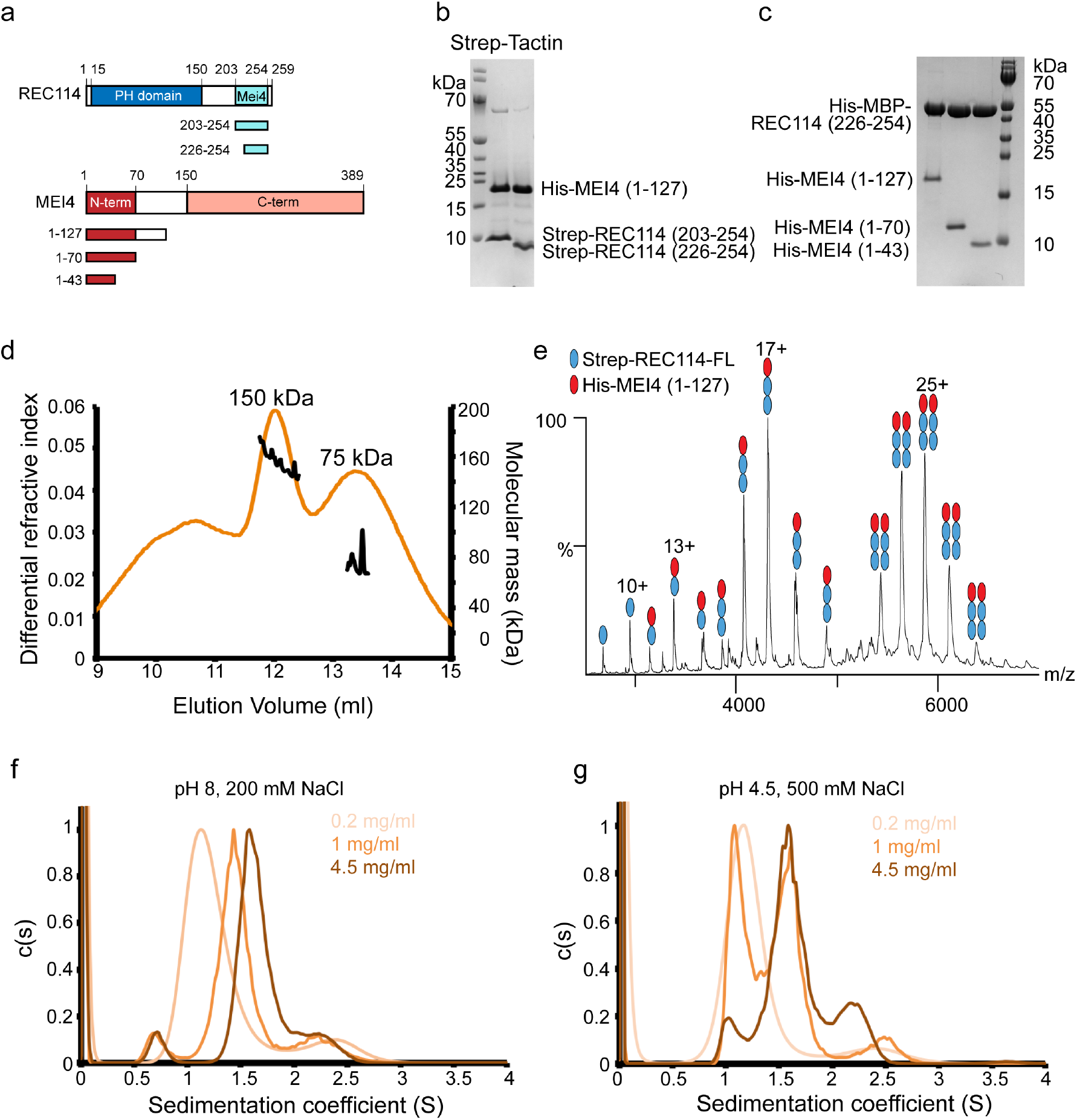
REC114-MEI4 complex stoichiometry characterization. **a**. Schematic representation of the domain structure of mouse REC114 (in red) and MEI4 (in blue) and the constructs used in this study. PH: Pleckstrin homology domain, MEI4: MEI4 binding region. **b**. SDS-PAGE analysis of the binding of full-length Strep-tagged REC114 variants (using a single Strep-tag (WSHPQFEK)) to co-expressed His-MEI4^1-127^ after purification on Strep-Tactin resin. **c**. SDS-PAGE analysis of the binding of His-MBP-REC114^226-254^ to co-expressed His-MEI4 variants after purification on Amylose resin. **d**. Molecular mass determination of the complex between the full-length Step-REC114 and His-MEI4^1-127^ by MALLS. The calculated molecular weight of REC114 is 29.5 kDa and of His-MEI4 is **e**. 17.5 kDa. The measured molecular mass of the two main peaks is 150 and 75 kDa corresponding to a REC114:MEI4 ratios 4:2 (153 kDa) and 2:1 (76.5 kDa), respectively. The injected sample was at 3 mg/ml. **f**. Native MS spectrum of co-purified Strep-REC114 and MEI4^1-127^. The main complexes are Strep-REC114:MEI4^1-127^ in ratios 2:1 and 4:2. There are also free Strep-REC114 and 1:1 Strep-REC114-MEI4 complex. **g**. Sedimentation velocity profiles of the REC114^226-254^-MEI4^1-43^ complex obtained at 280 nm at three concentrations (0.2 mg/ml, 1 mg/ml and 4.5 mg/ml) in a solution containing 20 mM Tris pH 8.0 and 200 mM NaCl. Expected sedimentation coefficient for the 2:1 complex is 1.13S and for 4:2 is 1.58S. The sedimentation coefficient of the main peak increases with increasing protein concentration: s=1.2S at 0.2 mg/ml, s=1.44S at 1 mg/ml and s=1.64S at 4 mg/ml. These results suggest a fast exchange between the two forms. **h**. Sedimentation velocity profiles of the REC114^226-254^-MEI4^1-43^ complex obtained at 280 nm at three concentrations (0.2 mg/ml, 1 mg/ml and 4.5 mg/ml) in a solution containing Sodium Acetate pH 4.5 and 500 mM NaCl. Expected sedimentation coefficient for the 2:1 complex is 1.13S and for the 4:2 complex is 1.58S. The sedimentation coefficient of the main peak is s=1.13S at 0.2 mg/ml and s=1.58S at 4 mg/ml. At 1 mg/ml both forms of REC114^226-254^-MEI4^1-43^ are present. At pH 4.5, a slow exchange between the two forms is observed.

To assess the stoichiometry of the complex between full-length Strep-REC114 and His-MEI4^1-127^ we first used multi angle laser light scattering (MALLS). The complex elutes in two major peaks (preceded a wider peak shoulder), with measured molecular weight of 150 and 75 kDa, respectively (Fig. 1d). These values correspond to the theoretical molecular masses of 4:2 (153 kDa) and 2:1 (76.5 kDa) stoichiometry of the REC114-MEI4 complex. Since the observed molecular weight is unstable in each peak, it is possible that these are mixtures of different oligomeric states. To obtain additional insight into the REC114-MEI4 complex architecture, we performed native mass spectrometry (MS). The analysed sample was a pool of the fractions of both main peaks from the gel filtration. In agreement with the MALLS results, we could detect complexes containing Strep-REC114 and MEI4^1-127^ in ratios 2:1 and 4:2. In addition, the MS data indicate presence of free Strep-REC114 and 1:1 Strep-REC114-MEI4^1-127^ complex (Fig. 1e). Finally, we analysed of the minimal REC114^226-254^-MEI4^1-43^ complex by analytical ultracentrifugation (AUC). The analysis confirmed the existence of predominant 2REC114-1MEI4 and 4REC114-2MEI4 complexes (Fig. 1f) (Supplementary Fig. 1). Measurements at different protein concentrations are consistent with a concentration-dependent dimerization of the 2REC114-1MEI4 heterotrimers, with fast exchange between the two states (Fig. 1f). We also noticed that the exchange between the two states was slower at low pH and high salt concentrating. In this condition, the two oligomeric states (2:1 and 4:2) could be detected with higher precision, as the exchange rate between them was slower (Fig. 1g). Together, these results show, that REC114 and MEI4 form predominantly a complex with a 2:1 stoichiometry that can further dimerize to 4:2 in response to increasing protein concentration.

### REC114-MEI4 complex structure prediction

We then attempted to determine the structure of the REC114-MEI4 minimal complex, but both crystallization and NMR analysis failed, likely due to the fast exchange between the different oligomeric states of the complex (Supplementary Fig. 2a). High accuracy predicted AlphaFold2 models of both REC114 and MEI4 are available in the EBI AlphaFold2 database (https://alphafold.ebi.ac.uk/) under the AF-Q9CWH4 and AF-Q8BRM6 accession numbers, respectively. Consistent, with previous experimental structures, REC114 is predicted to contain the PH domain in its N-terminus connected by a flexible linker to a short helical segment in its C-terminus that interacts with MEI4 (Supplementary Fig. 2b-d) (Kumar et al., 2018). MEI4 is predicted to contain a helical N-terminal segment (binding to REC114) connected flexibly to a larger helical domain (Supplementary Fig. 2e-g). It should be noted that in our hands, MEI4 constructs covering this helical domain could not be produced soluble in bacteria nor insect cells. AlphaFold2 (Jumper et al., 2021) predicts a very convincing model of the REC114-MEI4 2:1 heterotrimer (Fig. 2a-c, Supplementary Fig. 3a-d) that include contacts between the segments that we biochemically defined as minimal interacting regions (Fig. 1b,c). Predictions of 4:2 REC114-MEI4 complexes by AlphaFold2 were not convincing. In the predicted 2:1 model, the C-terminal helices α2 and α3 of two REC114 molecules wrap around the helix α1 of MEI4 (Fig. 2a-c). One of the REC114 protomers makes additional contact also with the MEI4 helix α2 (Fig. 2a-c, Supplementary Fig. 3d). Most of the predicted contacts are mediated by hydrophobic residues. On MEI4 α1, the interacting residues include a repetitive sequence 13-LALALAII-20 that form multiple contacts with REC114 F230, L231, F240, F243, V244 and V247 (Fig. 2c). While L13, A14, L17 and A18 interact with one REC114 protomer, L15, A16, I19 and I20 bind the other one (Fig. 2c). Within MEI4 helix α2, T33, L36, A37 and V40 pack against hydrophobic residues of one of the REC114 protomers, including L235, F240 and V244 (Supplementary Fig. 3d). Most residues predicted to interact are highly conserved in both proteins across vertebrates (Fig. 2d). Despite sequence divergence, equivalent models can also be predicted for other species, such as *A. thaliana, S. cerevisiae* and *S. pombe* (Supplementary Fig. 3e-g). To confirm that the residues predicted to mediate the interaction between REC114 and MEI4 are correct, we mutated REC114 residues F240, V244 and V247, respectively, to glutamates. In pull-down assays with the Strep-tagged full-length REC114, all three mutations F240E, V244E and V247E led to undetectable interaction with His-MBP-MEI4^1-43^ (Fig. 2e, lanes 10-12). When MEI4 α1 residues predicted as key for binding of REC114 were mutated and tested in pull-down assays with the Strep-tagged full-length REC114, the A16E mutation completely abolished the binding (Fig. 2f, lane 8). A significantly reduced binding was observed for L15E and a double mutant L13E,A14E (Fig. 2f, lanes 6-7). The mutations introduced into REC114 did not significantly alter the structure of these proteins as judged by gel filtration analysis (Supplementary Fig. 4a,b). His-MBP-MEI4^1-43^ tends to aggregate in absence of REC114, but only monomeric fractions of the WT and mutant versions were used for the pull-down experiment (Supplementary Figs. 4c-g). These mutagenesis results are in agreement with the AlphaFold2 prediction of the REC114-MEI4 complex structure.

**Figure 2.**
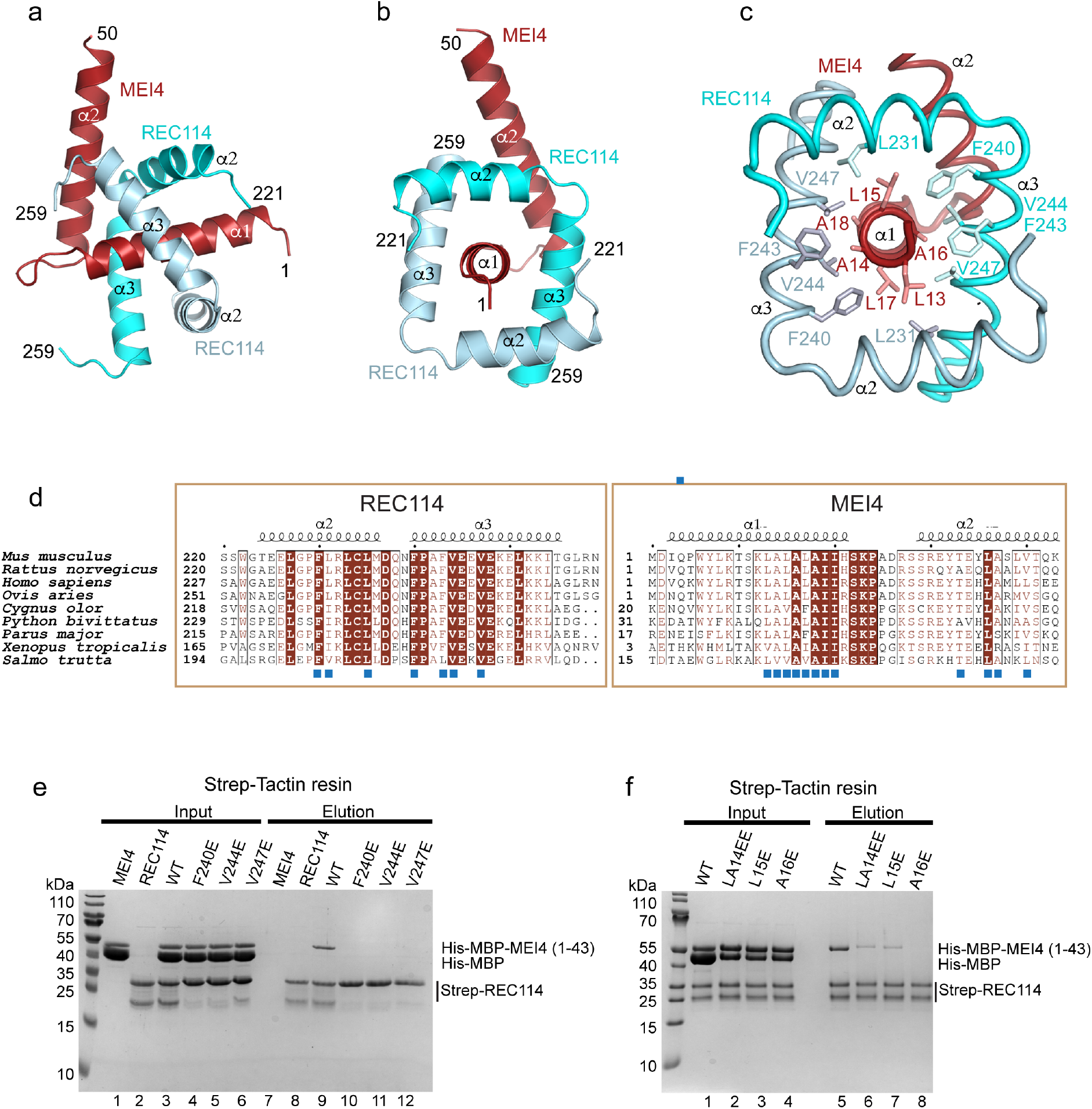
REC114-MEI4 complex structure prediction. **a**,**b** AlphaFold2 model of the REC114-MEI4 complex structure in 2:1 stoichiometry, including the REC114 C-terminus (residues 221-259) and the MEI4 N-terminus (residues 1-50). The secondary structures are labelled. The modelled complex structure coloured according to the AlphaFold2 per-residue estimate of confidence (pLDDT) and the predicted aligned error plot are shown in Supplementary Fig. 3a-c. **c**. Details of the predicted interactions between REC114 and MEI4. A sequence of hydrophobic residues of the MEI4 helix α1 is interacting with hydrophobic surfaces of the two molecules of REC114 wrapping around it. **d**. Sequence alignments of the interacting regions of REC114 and MEI4. Identical residues are in brown boxes. The residues involved in the interaction are shown as blue squares. **e**. Pull-down experiments of full-length Strep-tagged REC114 mutants indicated above the lanes with His-MBP-MEI4^1-43^. All proteins were first purified by affinity chromatography and gel filtration. A total of 0.9% of the input (lanes 1–6) and 1% of the eluates (lanes 7–12) were analyzed on 15% SDS-PAGE gels stained with coomassie brilliant blue. Control lanes 1-2 and 7-8 show inputs and elutions of REC114 and MEI4 alone, respectively. The MEI4 sample contains also free MBP tag. The REC114 sample also contains a degradation product, likely corresponding to its PH domain. **f**. Pull-down experiments of Strep-tagged REC114 with His-MBP-MEI4^1-43^ mutants indicated above the lanes. All proteins were first purified by affinity chromatography and gel filtration. A total of 0.9% of the input (lanes 1–4) and 1% of the eluates (lanes 5–8) were analyzed on 15% SDS-PAGE gels stained with coomassie brilliant blue. The MEI4 sample contains also free MBP tag. The REC114 sample also contains a degradation product, likely corresponding to its PH domain.

### REC114 forms homodimers

Since, in the AlpaFold2 model of the REC114-MEI4 complex, the two REC114 molecules interacting with MEI4 make mutual contacts (Fig. 2a-c) we next wanted to test whether the C-terminus of REC114 can dimerize also in absence of MEI4. MALLS analysis revealed that indeed His-MBP-REC114^226-254^ with theoretical molecular mass of 48kDa forms probable dimers (measured molecular weight of 86kDa) in solution (Fig. 3a). This was confirmed by native MS. We analysed the sample from the main gel filtration peak. MS data showed a clear dimer of His-MBP-REC114^226-254^ (Fig. 3b). Also AlphaFold2 predicts a REC114 dimer with high confidence (Fig. 3c-f, Supplementary Fig. 5a-c). The predicted structure of the REC114 dimer corresponds to that of REC114 in complex with MEI4. The contacts between the two REC114 protomers are located at the ends of both helices. In particular, the conserved L227 and L231 of α1 of one protomer are predicted to interact with area around I254 on α2 of the other protomer (Fig. 3e-f). In order to validate this model, we mutated residues I254 and L227, respectively, to aspartates and analysed their oligomeric state by MALLS. To accommodate potential extra contacts between the protomers predicted with low confidence the construct was slightly extended to residues 222-256. While the L227D eluted as a mixture of monomers and dimers, the I254D mutant was mostly monomeric (Fig. 3g). Mutations F240E, V244E and V247E which abolished the interaction between REC114 and MEI4 (Fig. 2e) did not have any impact on the dimerization of His-MBP-REC114^222-256^ being consistent with the AlphaFold2 model, according to which these residues do not play an important role in REC114 dimerization. In summary, these results show that the REC114 C-terminus is able to mediate REC114 homodimerization in absence of MEI4. But given the rather modest dimeric interface (Fig. 3 c-f), it is likely that MEI4 binding stabilises this complex.

**Figure 3.**
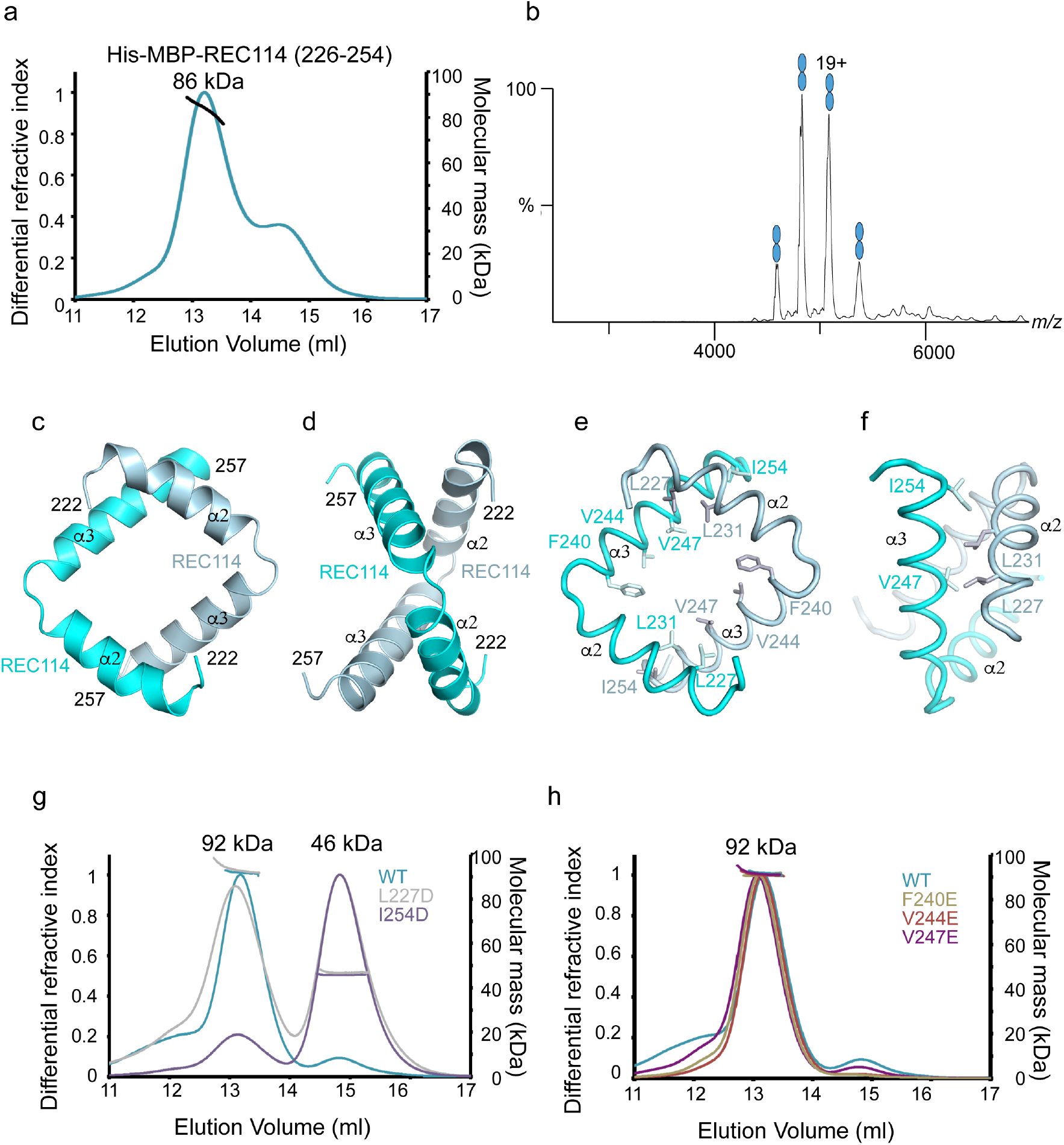
Characterization of the REC114 dimerization domain. **a**. Molecular mass determination of His-MBP-REC114^226-254^ by MALLS. The calculated molecular weight of His-MBP-REC114^226-254^ is 48 kDa. The measured molecular mass of the main peak is 86kDa which corresponds to dimeric His-MBP-REC114^226-254^ of 96 kDa. The injected sample was at 3mg/ml. **b**. Native MS spectrum of His-MBP-REC114^226-254^. This protein is a dimer. **c**,**d** AlphaFold2 model of the REC114 dimer including its C-terminus (residues 222-257). The secondary structures are labelled. The modelled complex structure coloured according to the AlphaFold2 per-residue estimate of confidence (pLDDT) and predicted aligned error plots are shown in Supplementary Fig. 5a-c. **e**,**f** Details of the predicted interactions between the two REC114 molecules formed by the extremities of the two helices. **g**. Comparison of molecular mass between WT and mutated versions of His-MBP-REC114^222-256^ by MALLS. L227D elutes partially as a monomer and I254D elutes mostly as a monomer. The injected samples were at 3 mg/ml. **h**. Comparison of molecular mass between WT and mutated versions of His-MBP-REC114^222-256^ by MALLS. All three tested REC114 mutants remain dimeric. The injected samples were at 3 mg/ml.

### IHO1 tetramerizes via a four-stranded coiled-coil

Mouse IHO1 is predicted to be largely disordered, except for its central region (110-240) predicted to be helical and possess a coiled-coil domain (Fig. 4a). We were not able to efficiently overexpress the full-length IHO1, but could express it in two fragments as His-MBP fusions: IHO1^1-281^ and IHO1^281-574^ (Fig. 4a). When IHO1^1-281^ was analysed by MALLS, a likely tetramer stoichiometry was observed (measured 120 kDa, calculated 126kDa for a tetramer) (Fig. 4b). The central region of IHO1 spanning residues 125-281 retained its tetrameric form (measured 67 kDa, calculated 71 kDa for a tetramer) (Fig. 4b). Finally, even the short fragment covering the C-terminal part of the IHO1 helical domain (residues 196-245) still forms a stable tetramer (measured 24 kDa, calculated 23 kDa for a tetramer) (Fig. 4b). The IHO1 AlphaFold2 model is available in the EBI AlphaFold2 database (https://alphafold.ebi.ac.uk/) under the AF-Q6PDM4 accession number, and confirms the presence of the long helix between residues 114-245 within otherwise disordered protein (Supplementary Fig. 6a,b). When a tetrameric stoichiometry is modelled, AlphaFold2 predicts with high confidence an IHO1 tetramer, based on a parallel four-stranded coiled-coil between residues 130 and 245 (Fig. 4c, Supplementary Fig. 6c). After residues 245 the prediction is less convincing. The sequence corresponding to the coiled-coil region is better conserved than the surrounding parts of IHO1 (Fig. 4d). Similar tetrameric models can also be modelled for IHO1 orthologs in other species, such as *A. thaliana, S. pombe* and *Sordaria macrospora* (Supplementary Fig. 7a-c). The model of *S. cerevisiae Mer2* is predicted with lower confidence (Supplementary Fig. 7d). While the N-terminal section of the coiled-coil contains a few irregularities in the heptad repeats, the *a* and *d* positions (the 1st and 4th positions in heptad repeats) are regularly spaced in the C-terminal part (residues 180-245) (Fig. 4d) being consistent with our MALLS results (Fig. 4b). These results thus clearly show that, similarly to *S. cerevisiae* Mer2, mouse IHO1 forms homotetramers (Claeys Bouuaert et al., 2021a; Rousova et al., 2021).

**Figure 4.**
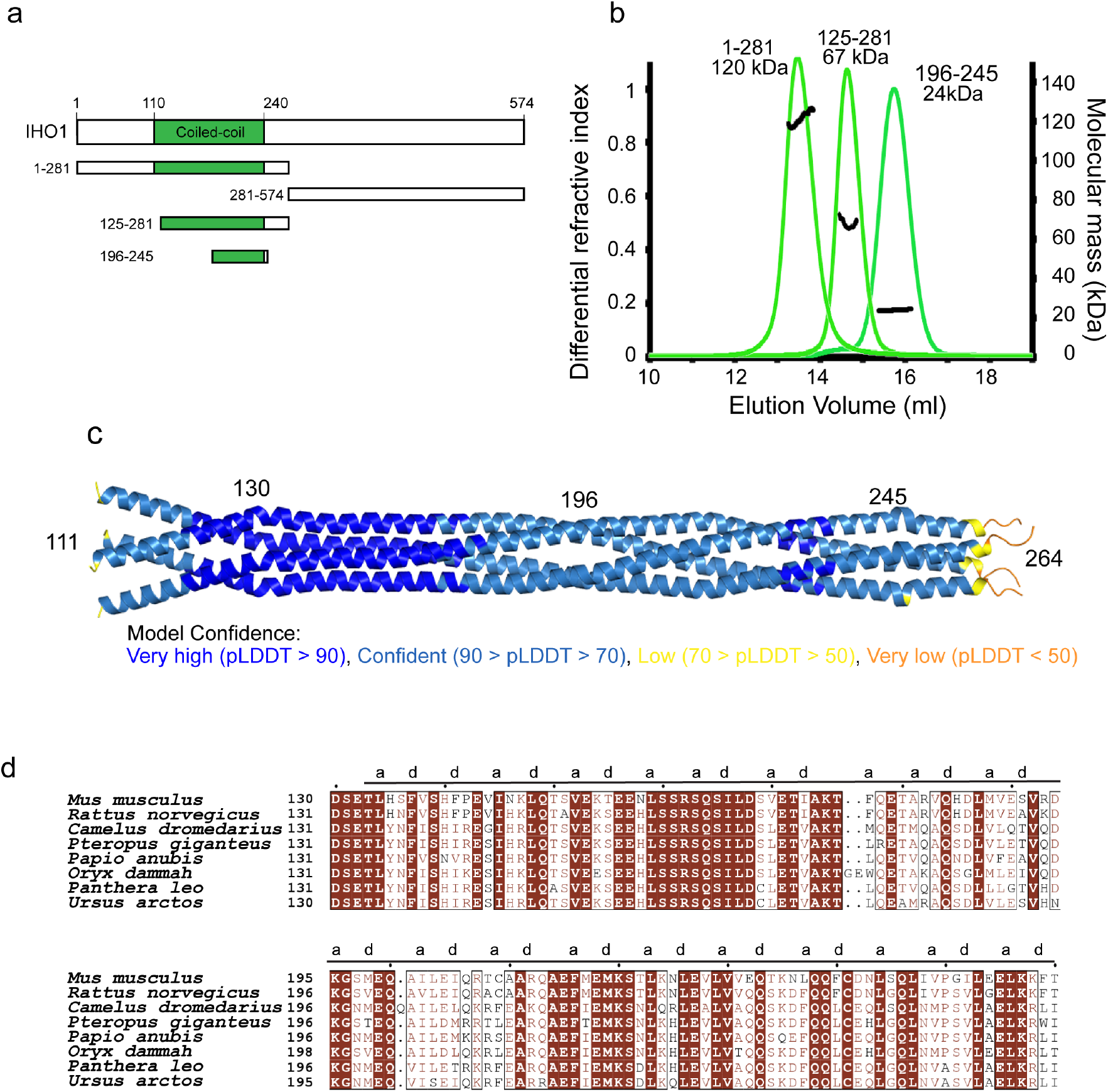
IHO1 forms tetramers. **a**. Schematic representation of the domain structure of mouse IHO1 and its constructs used in this study. **b**. Molecular mass determination of IHO1^1-281^, IHO1^125-281^ and IHO1^196-245^ by MALLS. The measured molecular mass of the main peak of each construct is 120kDa, 67kDa and 24kDa, respectively. These values correspond to calculated molecular weights of IHO1 tetramers (125 kDa, 72 Kda and 24 kD, respectively). The samples were injected at 5-10mg/ml. **c**. AlphaFold2 model of tetrameric IHO1^111-280^ coloured according to the AlphaFold2 per-residue estimate of confidence (pLDDT). The predicted aligned error plot for this model is shown in Supplementary Fig. 6c. **d**. Sequence alignments of the helical region of IHO1 (residues 130-260). Identical residues are in brown boxes. A and D positions of the heptad repeats are shown.

### IHO1 binds to the REC114 PH domain with its conserved N-terminus

IHO1 was previously shown to interact with REC114 in Y2H (Stanzione et al., 2016). We were thus interested in testing whether IHO1 can directly bind the REC114-MEI4^1-127^ in vitro. Interestingly, when the purified REC114-MEI4^1-127^ complex and His-MBP-IHO1^1-281^ were mixed and injected onto a Superose 6 gel filtration column, the three proteins co-eluted in separate peak, compared to His-MBP-IHO1^1-281^ and REC114-MEI4^1-127^ alone (Fig. 5a,b, Supplementary Fig. 8). Similarly, the three proteins co-elute together, when IHO1^1-281^ was fused to a His-Sumo tag (Supplementary Fig. 9). No complex formation was observed using IHO1^281-574^ (data not shown). Using Y2H, it has been shown, that the N-terminal region of the *Sordaria macrospora* IHO1 ortholog Asy2 (residues 1-156) is sufficient for REC114 binding (Tessé et al., 2017). Since this region bears a short motif with a distant similarity to the extreme N-terminus of mouse IHO1, we were interested in testing whether this motif is required for the interaction with REC114 in the mouse. Indeed, when the first conserved 23 residues were deleted from IHO1^1-281^ it could no longer interact with the REC114-MEI4^1-127^ complex (Fig. 5c,d, Supplementary Fig. 10). In agreement with these results, using isothermal titration calorimetry (ITC), we show that His-MBP-IHO1^1-281^ interacts with the REC114-MEI4^1-127^ complex with a dissociation constant (*K*d) of 26.9 µM (Fig. 5e), while no binding is observed for His-MBP-IHO1^24-281^ lacking the first 23 amino acids (Fig. 5f). In addition, we could show that the first 23 residues are sufficient for the interaction, as His-MBP-IHO1^1-23^ interacts with the REC114-MEI4^1-127^ complex with *K*d of 31.9 µM, being equivalent to the one of His-MBP-IHO1^1-281^ (Fig. 5g). Finally, ITC measurements revealed, that neither the REC114 C-terminus nor MEI4^1-127^ are required for the complex formation, since the REC114 PH domain (residues 1-159) binds His-MBP-IHO1^1-23^ with *K*d of 30.4 µM (Fig. 6a). These results provide evidence that IHO1 and REC114-MEI4 form a ternary complex, that mouse REC114 and IHO1 directly interact in vitro and the REC114 PH domain and the first 23 residues of IHO1 are the required regions for this interaction (Fig. 6b).

**Figure 5.**
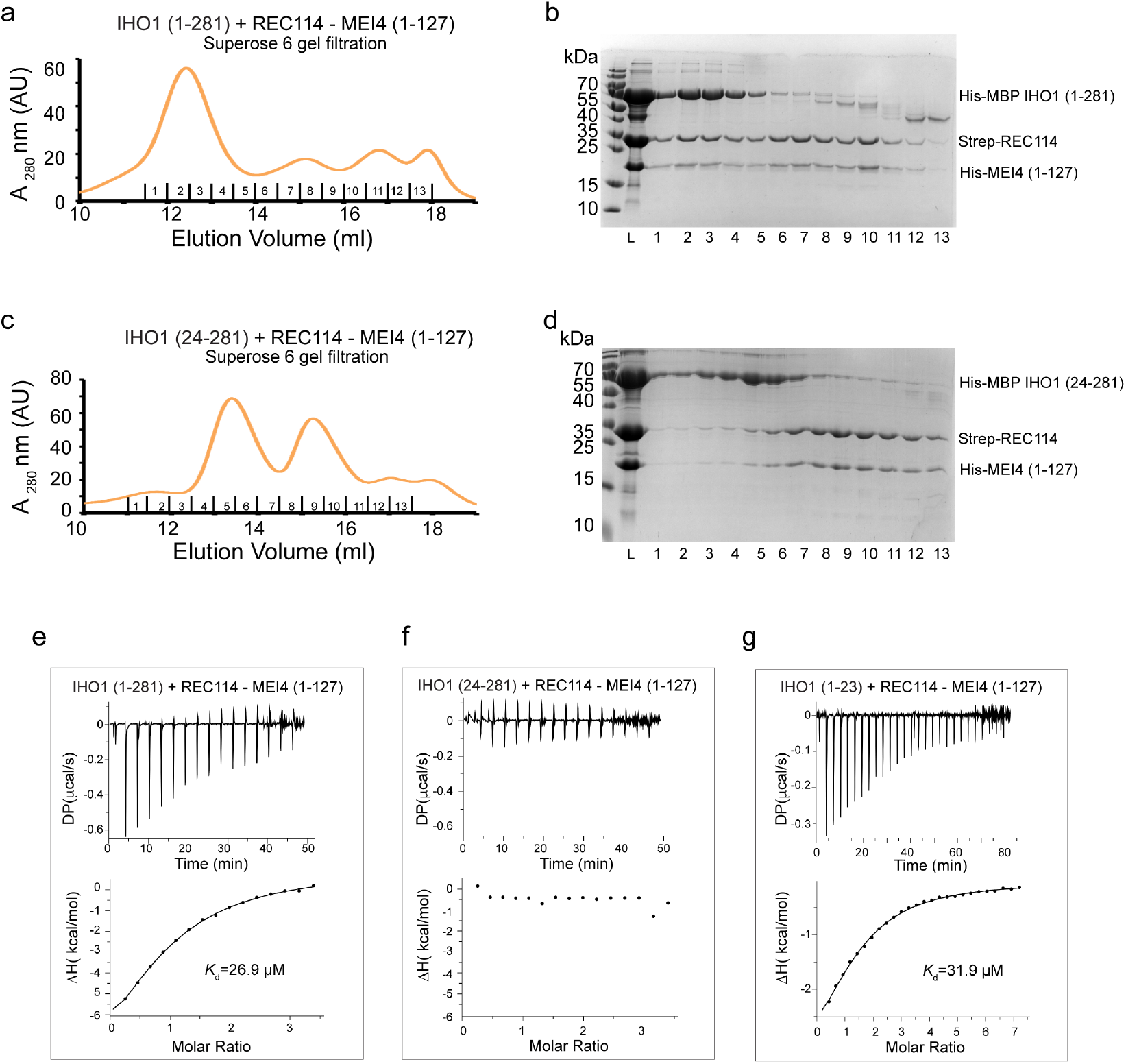
IHO1 forms a complex with REC114-MEI4. **a**. Superose 6 gel filtration elution profile of His-MBP-IHO1^1-281^ mixed with Strep-REC114-His-MEI4^1-127^. Overlay with individual elution profiles of His-MBP-IHO1^1-281^ and the Strep-REC114-His-MEI4^1-127^ complex and the corresponding SDS-PAGE analysis of the elution fractions are shown Supplementary Fig. 8. **b**. SDS-PAGE analysis of fraction 1-13 of the Superose 6 gel filtration elution profile shown in **a**. L indicates input sample loaded onto the column. **c**. Superose 6 gel filtration elution profile of His-MBP-IHO1^24-281^ mixed with Strep-REC114-His-MEI4^1-127^, showing lack of binding. Overlay with individual elution profiles of His-MBP-IHO1^24-281^ and the Strep-REC114-His-MEI4^1-127^ complex and the corresponding SDS-PAGE analysis of the elution fractions are shown Supplementary Fig. 10. **d**. SDS-PAGE analysis of fraction 1-13 of the Superose 6 gel filtration elution profile shown in **c**. L indicates input sample loaded onto the column. **e**. ITC measurement of the interaction affinity between of His-MBP-IHO1^1-281^ and Strep-REC114-His-MEI4^1-127^. **f**. ITC measurement of the interaction affinity between of His-MBP-IHO1^24-281^ and Strep-REC114-His-MEI4^1-127^. **g**. ITC measurement of the interaction affinity between of His-MBP-IHO1^1-23^ and Strep-REC114-His-MEI4^1-127^.

**Figure 6.**
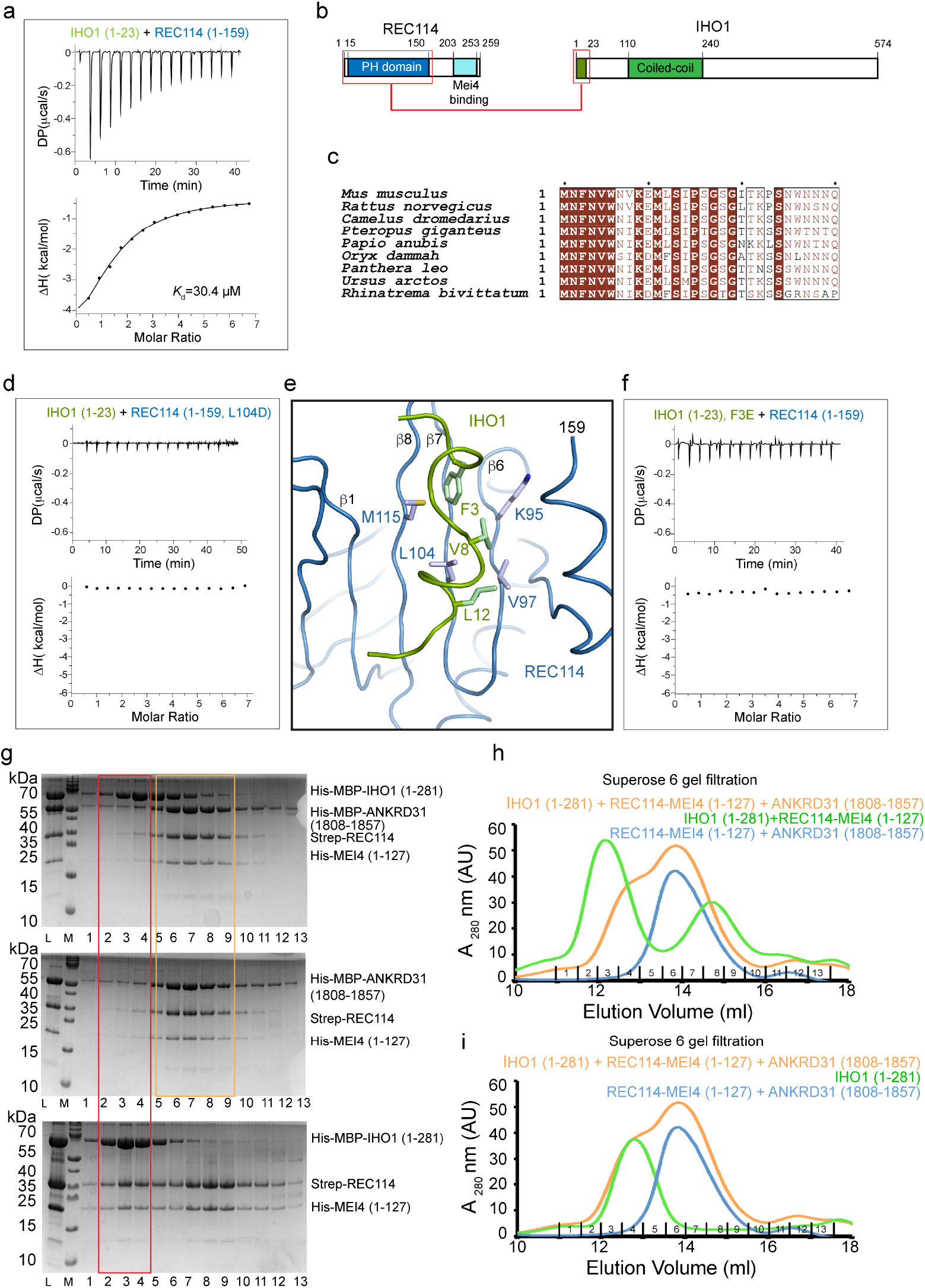
IHO1 interacts with the PH domain of REC114. **a**. ITC measurement of the interaction affinity between of His-MBP-IHO1^1-23^ and Strep-REC114^1-159^ (PH domain). **b**. Schematic representation of the interacting regions of REC114 and IHO1, highlighted with red boxes. **c**. Sequence alignments of the N-terminal sequence of IHO1. Identical residues are in brown boxes. Only first 30 residues are shown. **d**. ITC measurement of the interaction affinity between of His-MBP-IHO1^1-23^ and the L104D mutant of Strep-REC114^1-159^. **e**. AlphaFold2 model of the complex between the REC114 PH domain (blue) and the N-terminal peptide of IHO1 (residues 1-15, green). Hydrophobic IHO1 residues are predicted to interact with the REC114 β-sheet formed of strands β1 and β6-β8. The secondary structures are labelled. The modelled complex structure coloured according to the AlphaFold2 per-residue estimate of confidence (pLDDT) and predicted aligned error plot are shown in Supplementary Fig. 11a-b. **f**. ITC measurement of the interaction affinity between of the F3E mutant of His-MBP-IHO1^1-23^ and Strep-REC114^1-159^. **g**. SDS-PAGE analysis of fractions 1-13 of Superose 6 gel filtration elution profiles of His-MBP-IHO1^1-281^ mixed with His-MBP-ANKRD31^1808-1857^, Strep-REC114 and His-MEI4^1-127^ (upper panel); mixture of His-MBP-ANKRD31^1808-1857^ with Strep-REC114-His-MEI4^1-127^ (middle panel) and His-MBP-IHO1^1-281^ mixed with Strep-REC114 and His-MEI4^1-127^ (lower panel). L indicates input sample loaded onto the column. The red frame highlights the complex of IHO1 bound to REC114 and MEI4 (lower panel) and the lack of this complex in presence of ANKRD31 (upper panel). Orange square shows equivalent elution of ANKRD31, REC114 and MEI4 in presence or absence of IHO1. **h**. Superose 6 gel filtration elution profiles of His-MBP-IHO1^1-281^ mixed with His-MBP-ANKRD31^1808-1857^, Strep-REC114 and His-MEI4^1-127^ (orange); His-MBP-ANKRD31^1808-1857^, Strep-REC114 and His-MEI4^1-127^ (blue) and His-MBP-IHO1^1-281^ mixed with Strep-REC114-His-MEI4^1-127^ (green). Corresponding SDS-PAGE gel analysis is shown in **g**. **i**. Superose 6 gel filtration elution profiles of His-MBP-IHO1^1-281^ mixed with His-MBP-ANKRD31^1808-1857^, Strep-REC114 and His-MEI4^1-127^ (orange); His-MBP-ANKRD31^1808-1857^, Strep-REC114 and His-MEI4^1-127^ (blue) and His-MBP-IHO1^1-281^ (green). Corresponding SDS-PAGE gel analysis is shown in Supplementary Fig. 12.

### IHO1, ANKRD31 and TOPOVIBL bind to the same REC114 surface

The REC114-binding region of IHO1 is very well conserved among mammals (Fig. 6c) and some sequence similarities have been reported across species up to *S. cerevisiae* (Tessé et al., 2017). The PH domain has previously been shown to interact via its β-sheet formed of strands β1, β2 and β6-β8 in a mutually exclusive way with hydrophobic and aromatic residues of short motifs of TOPOVIBL (Nore et al., 2022) and ANKRD31 (Boekhout et al., 2019). A mutation of L104 on its β-strand β7 abolished the binding to both ANKRD31 (Boekhout et al., 2019) and to TOPOVIBL (Nore et al., 2022). We could show by ITC that the L104D mutation prevents also the interaction between REC114 and IHO1 suggesting that IHO1 uses this β-sheet surface for the interaction with REC114 too (Fig. 6d). The L104D mutation does not impact the REC114 PH domain structure (Nore et al., 2022). AlphaFold2 models the possible structure of the complex between the REC114 PH domain and the IHO1^1-30^ peptide with a low confidence (Fig. 6e, Supplementary Fig. 11a,b). However, the predicted aligned error plot shows certain degree of confidence about the mutual positions of the two proteins (Supplementary Fig. 11b). Since, in this model the IHO1 peptide is predicted to interact with L104 that is indeed required for the interaction, we also mutated the F3 of IHO1, predicted to insert into the cavity that can also accommodate W562 of TOPOVIBL (Nore et al., 2022) and W1842 of ANKRD31 (Boekhout et al., 2019), both essential for binding to REC114. ITC measurement confirmed that the F3E mutation in His-MBP-IHO1^1-23^ abolished the interaction with REC114 (Fig. 6f). While the AlphaFold2 model is not of sufficient quality to be certain of positions of individual residues of IHO1, our biochemical results demonstrate, that IHO1 uses an equivalent surface on REC114 as do TOPOVIBL and ANKRD31 (Supplementary Fig. 11c-e).

To further test whether IHO1 binding to REC114 is mutually exclusive with the binding of ANKRD31, we wanted to test whether quaternary complexes including these proteins can be formed on Superose 6 gel filtration. To this end we prepared a complex between Strep-REC114, His-MEI4^1-127^ and MBP-ANKRD31^1808-1857^ that elutes as a distinct single peak (Fig. 6g, middle panel and Fig. 6h), showing that REC114 can bind ANKRD31 and MEI4 at the same time. We then mixed MBP-IHO1^1-281^ with this REC114-MEI4-ANKRD31 complex. As expected, in presence of ANKRD31^1808-1857^ IHO1 does not bind REC114-MEI4 anymore, and those elute only in complex with ANKRD31^1808-1857^ (Fig. 6g, top panel and Fig. 6h). The elution profile of IHO1 in mixture with the REC114-MEI4-ANKRD31 complex is essentially the same as when IHO1 is injected on its own (Fig. 6i, Supplementary Fig. 12). In absence of ANKRD31^1808-1857^, IHO1 forms a complex with REC114 and MEI4, as shown previously in Fig. 4b (Fig. 6g, bottom panel and Fig. 6h). These results show that a REC114 does not interact at the same time with IHO1 and ANKRD31. When TOPOVIBL^452-579^ is mixed with the complex of Strep-REC114-His-MEI4^1-127^ the resulting complex elutes on Superose 6 in a large peak overlapping with the elution volume of MBP-IHO1^1-281^, making an equivalent experiment with TOPOVIBL^452-579^ less interpretable. It can, however, be noted that the His-MBP-IHO1^1-281^ elution peak is not shifted when mixed with Strep-REC114-His-MEI4^1-127^ in presence of TOPOVIBL^452-579^ indicating a probable lack of complex formation (Supplementary Fig. 13). These results thus further show that the binding of IHO1 to REC114 is incompatible with the binging of ANKRD31 or TOPOVIBL.

## Discussion

REC114, MEI4 and IHO are known to be essential for DSB formation and based on their co-localization and Y2H assays they were proposed to function as a complex (Pre-DSB recombinosome), that regulates the TOPOVIL catalytic complex activity (Dereli et al., 2021; Kumar et al., 2018, 2015, 2010; Stanzione et al., 2016). The interaction network as well as molecular architecture of such a complex remain unclear.

Mouse REC114 possesses a PH domain in its N-terminus that forms mutually exclusive interactions with ANKRD31 (Boekhout et al., 2019) and the TOPOVIBL subunit of the catalytic complex forming DSBs (Nore et al., 2022). In addition, its C-terminal region (residues 203-259) interacts with MEI4^1-127^ in pull-down assays and gel filtration (Kumar et al., 2018). Similarly, in *S. cerevisiae* these two proteins interact with equivalent regions, likely with a 2:1 stoichiometry (Claeys Bouuaert et al., 2021a). We could now show that, in the mouse, the two interacting regions can be further truncated to REC114^226-254^ and MEI4^1-43^. Our attempts to structurally characterise this complex were not successful. In particular, different approaches to analyse the complex by NMR indicated that the complex has a molecular mass higher than expected and is subject to fast exchange between different molecular states. Using native MS we confirmed this hypothesis. Indeed, REC114 and MEI4^1-127^ form a mixture of 2:1 and 4:2 complexes. In addition, analytical ultracentrifugation showed that even the minimal complex REC114^226-254^-MEI4^1-43^ forms 2:1 and 4:2 complex with fast exchange at pH 8 and slower exchange at pH 4.5. AlphaFold2 predicts with high confidence the structure of the 2:1 REC114:MEI4 complex with contacts between the two short segments we identified in vitro. Mutations of REC114 and MEI4 residues predicted to be key for the interaction indeed abolished the binding in pull-down assays, providing further evidence for correctness of the model. The interacting residues are conserved in mammals and equivalent predication can also be made for more distantly related species up to *S. cerevisiae*, where sequence conservation of the two proteins is less obvious. Overall, even in absence of experimental structure, these results support the 2:1 stoichiometry of the complex and identified residues essential for the binding.

REC114^226-254^ forms homodimers as judged by MALLS and native MS even in absence of MEI4. Similarly, the *S. cerevisiae* Rec114 forms Mei4-independent homodimers (Claeys Bouuaert et al., 2021a). AlphaFold2 predicted structure of the REC114 dimer corresponds to the REC114 dimer in complex with MEI4 and we confirmed the residues essential for the homodimerization by mutagenesis. AlphaFold2 does not predict a convincing model for the 4:2 complex and atomic details of the dimerization of the REC114-MEI4 heterotrimers thus remain unknown. It should be noted, that we were not able to produce MEI4 containing also its C-terminal domain, which might provide further dimerization contacts.

REC114 thus forms homodimers, and it is likely that these are further stabilised by MEI4. We recently reported that REC114-MEI4^1-127^ forms a stable complex with TOPOVIBL^452-579^ (Nore et al., 2022). Here we show that REC114-MEI4^1-127^ forms a stable complex with ANKRD31^1808-1857^. One of the functions of the REC114-MEI4 might to contribute to dimerization of ANKRD31 and TOPOVIBL. Indeed, the AlphaFold2 model of the mouse SPO11-TOPOVIBL complex indicates that the putative TOPOVIBL ATP-binding site required for its ATP-mediated dimerization is degenerated (Nore et al., 2022). Hence, the interaction with REC114-MEI4 might provide additional means of dimerization of the catalytic complex.

IHO1 was shown to interact with REC114 in Y2H assay but direct interaction between the two proteins could not be detected. In *S. cerevisiae*, Rec114-Mei4 and Mer2 (IHO1 ortholog) were proposed not to form a complex, but rather to create common DNA-based condensates. Here, we demonstrate by gel-filtration and ITC measurement that in the mouse, IHO1 and REC114-MEI4 form a stable complex. This interaction requires a highly conserved N-terminus of IHO1 and the PH domain of REC114 and the *K*d of the interaction measured by ITC is 30.4 µM. The AlphaFold2 model of the complex is not predicted with high confidence, nevertheless, it guided us to identify conserved residues of REC114 and IHO1 that are indeed required for the interaction.

We could also show by MALLS that mouse IHO1 forms tetramers with its central helical region. This has already been shown for the yeast Mer2(Claeys Bouuaert et al., 2021a; Rousova et al., 2021). AlphaFold2 predicts the tetramerization region to be a long four-stranded parallel coiled-coil for mouse and other species. Given that REC114 and MEI4 form 2:1 and 4:2 complexes and each PH domain of REC114 possess a binding site for one N-terminal peptide of IHO1, it is tempting to speculate that the REC114-MEI4-IHO1 complex might be formed of 4 IHO1, 4 REC114 and 2 MEI4 molecules (Fig. 7). Native MS and MALLS characterization of such a complex did not yield convincing stoichiometry values, likely due to the fast exchange and low binding affinity. Importantly, the IHO1 binding requires the same REC114 residue L104, that is positioned centrally in its β-sheet and that interacts with hydrophobic and aromatic residues of short motifs of TOPOVIBL (Nore et al., 2022) and ANKRD31 (Boekhout et al., 2019), indicating a shared interacting surface and hence mutually exclusive binding of these proteins. The binding of ANKRD31 and TOPOVIBL to REC114 is indeed incompatible (Nore et al., 2022). We could show, using size exclusion chromatography, that in presence of ANKRD31^1808-1857^, IHO1 does not bind REC114-MEI4 anymore. The same is probably true for TOPOVIBL, since no peak shift of IHO1 is observed when REC114-MEI4 is first mixed with TOPOVIBL^452-579^. The binding affinity of ANKRD31 for REC114 is not known, since the ANKRD31 peptide aggregates in absence of REC114, but given the large interaction interface on REC114 it is likely higher than that of TOPOVIB (*K*d =3 µM (Nore et al., 2022)). IHO1 thus binds to REC114 with lower affinity than TOPOIVBL and ANKRD31, even though within the larger IHO1-REC114-MEI4 complex containing 4 REC114-IHO1 interfaces, the affinity might be higher. The affinity might as well be modified by posttranslational modifications or other protein or DNA factors. Indeed, in yeast, recruitment of Rec114 and Mei4 to chromosomal axis depends on Mer2 phosphorylation by Cdk (Henderson et al., 2006; Panizza et al., 2011; Sasanuma et al., 2008). MEI4 also co-immunoprecipitates with IHO1 even in cells lacking REC114, indicated other potential contacts between these proteins (Kumar et al., 2018). The role of REC114 and its multiple interactions in DSB formation regulation thus seems to be more complex. It is interesting to note that REC114 association to meiotic chromosome axis in mice depends on IHO1 in most of the genome expect for subtelomeric regions of some chromosomes where REC114 binding is independent of IHO1 (Papanikos et al., 2019). These properties may be due to the formation of alternative complexes. How IHO1, TOPOVIBL and ANKRD31 succeed each other on the surface of REC114 in vivo remains to be established.

**Figure 7.**
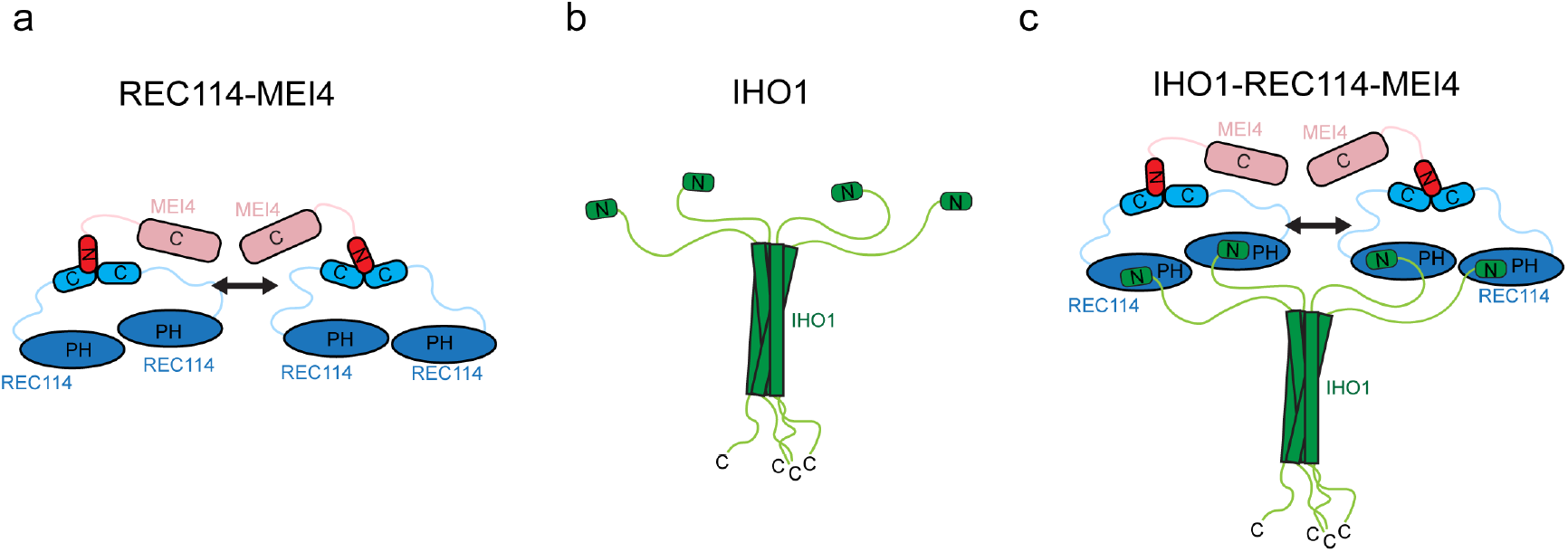
Model of the IHO1-REC114-MEI4 complex. **a**. Model summarising the current knowledge on the REC114-MEI4 complex architecture. REC114-MEI4 form 2:1 and 4:2 complexes through the C-termini of REC114 and N-terminus of MEI4. The arrow indicates potential dimerization of the REC114-MEI4 heterotrimers. **b**. Model summarising the current knowledge on IHO1 domain architecture. IHO1 forms tetramers and interacts with REC114 with its conserved N-terminus. **c**. Proposed model of the 4:4:2 IHO1-REC114-MEI4 complex. Tetramer of IHO1 interacts with 4 molecules of REC114 binding 2 molecules of MEI4. The arrow indicates potential dimerization of the REC114-MEI4 heterotrimers.

## Methods

### Protein Expression and Purification

All IHO1 constructs were expressed as His-MBP-tag fusions from the pETM41 vector (EMBL), or as a His-Sumo-tag fusion from pETM11 vector (EMBL) in *E. coli* BL21-Gold cells (DE3, Agilent). The protein was first purified on amylose resin (NEB) in a buffer containing 20 mM Tris pH 8, 100 mM NaCl, 2 mM β-mercaptoethanol, 5% glycerol and 1mM EDTA. To prepare an untagged protein, the His-MBP tag was cleaved off by TEV protease and the protein was further purified by a passage through a Ni^2+-^Chelating Sepharose (GE Healthcare). IHO1 was then purified by a gel filtration on Superdex 200 (GE Healthcare).

REC114^226-254^, REC114^222-257^ and its mutants were expressed in *E. coli* BL21-Gold cells (DE3, Agilent) from pETM41 vector (EMBL) as His-MBP-tag fusions. REC114^1-159^ and its mutant were cloned into pRSFDuet-1 vector as Strep-tag fusions. The proteins were, respectively, purified by affinity chromatography (amylose resin (NEB) or Strep-Tactin XT resin (IBA)) and by Superdex 200 size-exclusion chromatography in a buffer containing 20mM Tris pH 8, 100 mM NaCl and 2 mM β-mercaptoethanol.

### Protein complex reconstitution

Strep-Full-length REC114 and His-MEI4^1-127^ were co-expressed from pRSFDuet-1 (Novagen) and pProEXHTb (Invitrogen) vectors, respectively, in *E. coli* BL21Gold (DE3) cells. The REC114-MEI4 complex was purified using a Strep-Tactin XT resin (IBA) and by a gel filtration on Superdex 200 (GE Healthcare). For complex formation with IHO1, the REC114-MEI4 complex was first mixed with the purified His-MBP-IHO1 (or His-Sumo-IHO1) in a molar ratio of 2:1. The sample was then applied onto a Superose 6 gel filtration column (GE Healthcare) in 20 mM Tris pH 8, 100 mM NaCl, 5% glycerol and 2 mM β-mercaptoethanol.

His-MBP-REC114^226-254^ and His-MEI4^1-43^ were, co-expressed from pETM41 (EMBL) and pProEXHTb (Invitrogen) vectors, respectively, in *E. coli* BL21-Gold cells (DE3, Agilent) and were first purified by affinity chromatography using the amylose resin (NEB). For untagged complex, the His-MBP and His tags were removed by TEV protease and a passage through a Ni^2+^chelating Sepharose (GE Healthcare). The final purification step was size-exclusion chromatography on Superdex 200 size exclusion chromatography (GE Healthcare).

ANKRD31^1808–1857^ was cloned as His-MBP fusion into pETM41 (EMBL). The Strep-FL-REC114-His-MEI4^1-127^ complex and ANKRD31^1808–1857^ were individually expressed in *E. coli* BL21Gold (DE3) cells. Following cell disruption, supernatants containing soluble Strep-FL-REC114-His-MEI4^1-127^and ANKRD31^1808–1857^ were mixed. The ternary complex was further purified by Strep-Tactin XT resin (IBA) and Superose 6 size exclusion chromatography (GE Healthcare).

TOPOVIBL^452-579^ was cloned as a His-tag fusion into pProEXHTb (Invitrogen). REC114-MEI4 complex and TOPOVIBL^452-579^ were individually expressed in *E. coli* BL21Gold (DE3) cells and purified, respectively, on Strep-Tactin XT resin (IBA) or Ni^2+-^Chelating Sepharose (GE Healthcare). Proteins were mixed in a molar ratio of 2:1 and further purified by gel filtration on Superose 6 (GE, Healthcare).

### Pull-down Assays

MEI4 constructs were cloned into pProEXHTb (Invitrogen) as a His-tag fusions and REC114^226-254^ into pETM41 vector as a His-MBP fusion. His-MBP-REC114^226-254^ was co-expressed with the different constructs of MEI4 in *E. coli* BL21-Gold (DE3) cells (Agilent). Following cell disruption, the REC114-MEI4 protein-containing supernatants were loaded onto an Amylose resin (NEB). After extensive washing with a buffer containing 20mM Tris pH 8, 100 mM NaCl, 5% glycerol and 2 mM β-mercaptoethanol. Bound proteins were eluted by addition of 10mM maltose, and analysed on 15% SDS-PAGE. For analysis of the interaction between REC114 constructs and MEI4, REC114 variants were cloned as N-terminal Strep-tag fusions into pRSFDuet-1 (Novagen) and co-expressed with MEI4^1-127^ that was cloned as a His-tag fusion into pProEXHTb (Invitrogen). Supernatants were applied to Strep-Tactin XT resin (IBA) that was extensively washed with 20mM Tris pH 8, 100 mM NaCl, 5% glycerol and 2 mM β-mercaptoethanol and the bound proteins were eluted with addition of 50mM of D-Biotin.

To analyse the impact of REC114 and MEI4 mutants, full-length REC114 and mutants were cloned as Strep-tag fusions into pRSFDuet-1 (Novagen). MEI4^1-43^ and its mutants were cloned as His-MBP fusions in pETM41. Proteins were expressed individually in *E. coli* BL21Gold (DE3) cells. REC114 and MEI4 variants were purified by affinity chromatography (Strep-Tactin XT resin (IBA) and Ni^2+-^Chelating Sepharose, GE Healthcare, respectively) and gel filtration on Superdex 200 (GE, Healthcare). REC114 and MEI4 were mixed and loaded onto Strep-Tactin XT (IBA) resin columns. Columns were extensively washed with 20mM Tris pH 8, 100 mM NaCl, 5% glycerol and 2 mM β-mercaptoethanol, and bound proteins were eluted by addition of 50mM of D-Biotin and analysed by 15% SDS-PAGE.

### Isothermal Titration Calorimetry (ITC)

ITC experiments were performed at 25°C using an ITC200 microcalorimeter (MicroCal). Experiments included one 0.5µl injection and 15-25 injections of 1.5-2.5µL of 0.5-1mM His-MBP-IHO1 (IHO1^1-281^, IHO1^1-23^or IHO1^1-23, F3E^) into the sample cell that contained 30µM Strep-REC114-His-MEI4^1-127^ (or Strep-REC114^1-159^ or Strep-REC114^1-159, L104D^) in 20 mM Tris pH 8.0, 100 mM NaCl and 2 mM β-mercaptoethanol. The initial data point was deleted from the data sets. Binding isotherms were fitted with a one-site binding model by nonlinear regression using the MicroCal PEAQ-ITC Analysis software.

### Size-exclusion chromatography coupled to multi-angle laser light scattering

SEC-MALLS experiments were performed on high-performance liquid chromatography (HPLC) system (Shimadzu, Kyoto, Japan), consisting of a DGU-20 AD degasser, an LC-20 AD pump, a SIL20-ACHT autosampler, an XL-Therm column oven (WynSep, Sainte Foy d’Aigrefeuille, France), a CBM-20A communication interface, an SPD-M20A UV-visible detector, a miniDAWN TREOS static light scattering detector, a DynaPro NanoStar dynamic light scattering detector, and an Optilab rEX refractive index detector (Wyatt, Santa Barbara, USA). The samples were stored at 4°C, and a volume of 50 – 100 µL was injected on a Superdex 200 size exclusion column equilibrated with 20 mM Tris pH 8.0 and 100 mM NaCl, filtered at 0.1 µm, at a flow rate of 0.5 mL.min −1. The analysis of the data was done with the software ASTRA v5 (Wyatt, Santa Barbara, USA).

### Analytical ultracentrifugation

Sedimentation velocity experiments were performed at 42 000 rpm and 10°C, on a Beckman XLI analytical ultracentrifuge using a AN-50 Ti rotor (Beckman Coulter, Brea, USA) and double-sector cells with optical path lengths of 12 and 3 mm equipped with sapphire windows (Nanolytics, Potsdam, DE). The reference were samples buffer, for pH 8.0 samples: 200 mM NaCl, 20 mM Tris pH 8.0 and for pH 4.5 samples: 500 mM NaCl, 20 mM Sodium Acetate pH 4.5. Measurements were made on 0.2, 1 and 4.5 mg.ml–1 DdrC using absorbance at 280 nm and interference optics. Data were processed with the REDATE software (https://www.utsouthwestern.edu/labs/mbr/software/) and the parameters were determined with SEDNTERP and SEDFIT(Schuck, 2000). Analysis of sedimentation coefficients and molecular weights were performed using SEDFIT (Schuck, 2000) and GUSSI (Brautigam, 2015).

### Liquid chromatography/electrospray-ionization mass spectrometry (LC/ESI-TOF-MS)

To assess the mass of the different proteins, a 6210 TOF mass spectrometer coupled to a HPLC system (1100 series, Agilent Technologies) was used. The mass spectrometer was calibrated with tuning mix (ESI-L, Agilent Technologies). The following instrumental settings were used: gas temperature (nitrogen) 300 ◦C, drying gas (nitrogen) 7 L/min, nebulizer gas (nitrogen) 10 psi, Vcap 4 kV, fragmentor 250 V, skimmer 60 V, Vpp (octopole RF) 250 V. The HPLC mobile phases were prepared with HPLC grade solvents. Mobile phase A composition was: H2O 95%, ACN 5%, TFA 0.03%. Mobile phase B composition was: ACN 95%, H2O 5%, TFA 0.03%. Each protein was diluted to 5 uM using mobile phase A. 4 µl of each sample (20 pmol) were injected into HPLC system MS analysis and were first desalted on-line for 3 min with 100% of mobile phase A (flow rate of 50 µl/ min), using a C8 reverse phase micro-column (Zorbax 300SB-C8, 5 µm, 5 × 0.3 mm, Agilent Technologies). The sample was then eluted with 70% of mobile phase B (flow rate of 50 µl/ min) and MS spectra were acquired in the positive ion mode in the 300–3000 m/z range(Boeri Erba et al., 2018). Data were processed using MassHunter software (v. B.02.00, Agilent Technologies) and GPMAW software (v. 7.00b2, Lighthouse Data, Denmark).

### Native mass spectrometry

The samples were analyzed by native mass spectrometry (Boeri Erba et al., 2020, 2018; Puglisi et al., 2020). Protein ions were generated using a nanoflow electrospray (nano-ESI) source. Nanoflow platinum-coated borosilicate electrospray capillaries were bought from Thermo Electron SAS (Courtaboeuf, France). MS analyses were carried out on a quadrupole time-of-flight mass spectrometer (Q-TOF Ultima, Waters Corporation, Manchester, U.K.). The instrument was modified for the detection of high masses (Sobott et al., 2002; van den Heuvel et al., 2006). The following instrumental parameters were used: capillary voltage = 1.2–1.3 kV, cone potential = 40 V, RF lens-1 potential = 40 V, RF lens-2 potential = 1 V, aperture-1 potential = 0 V, collision energy = 30–140 V, and microchannel plate (MCP) = 1900 V. All mass spectra were calibrated externally using a solution of cesium iodide (6 mg/mL in 50% isopropanol) and were processed using the Masslynx 4.0 software (Waters Corporation, Manchester, U.K.) Massign software package (Morgner and Robinson, 2012) and UniDec (Marty et al., 2015).

## Acknowledgements

JK and BdM were funded ANR Topobreaks (ANR-18-CE11-0024-01). BdM was funded by ERC (European Research Council (ERC) Executive Agency under the European Union’s Horizon 2020 research and innovation programme (Grant Agreement no. 883605)). Ariadna B. Juarez-Martinez was supported by the Labex GRAL (Grenoble Alliance for Integrated Structural Cell Biology) (ANR-10-LABX-49-01) and the People Programme (Marie Curie Actions) of the European Union’s Seventh Framework Programme (FP7/2007-2013) under REA grant agreement PCOFUND-GA-2013-609102, through the PRESTIGE programme coordinated by Campus France. Financial support from the Centre National de la Recherche Scientifique (IR-RMN-THC Fr3050) is gratefully acknowledged. IBS acknowledges integration into the Interdisciplinary Research Institute of Grenoble (IRIG, CEA). This work used the platforms of the Grenoble Instruct-ERIC center (ISBG; UAR 3518 CNRS-CEA-UGA-EMBL) within the Grenoble Partnership for Structural Biology (PSB), supported by FRISBI (ANR-10-INBS-0005-02) and GRAL, financed within the University Grenoble Alpes graduate school (Ecoles Universitaires de Recherche) CBH-EUR-GS (ANR-17-EURE-0003). We thank Caroline Mas for assistance with ITC. We thank Adrien Favier and Cameron D. Mackereth for initial NMR analysis. We thank Henri-Marc Bourbon for discussions on the AlphaFold2 models.

## References

Acquaviva, L., Boekhout, M., Karasu, M.E., Brick, K., Pratto, F., Li, T., van Overbeek, M., Kauppi, L., Camerini-Otero, R.D., Jasin, M., Keeney, S. (2020). Ensuring meiotic DNA break formation in the mouse pseudoautosomal region. Nature 582, 426–431.

Arora, C., Kee, K., Maleki, S., Keeney, S. (2004). Antiviral protein Ski8 is a direct partner of Spo11 in meiotic DNA break formation, independent of its cytoplasmic role in RNA metabolism. Mol. Cell 13, 549–559.

Baudat, F., Imai, Y., De Massy, B. (2013). Meiotic recombination in mammals: Localization and regulation. Nat. Rev. Genet. 14, 794–806.

Baudat, F., Manova, K., Yuen, J.P., Jasin, M., Keeney, S. (2000). Chromosome synapsis defects and sexually dimorphic meiotic progression in mice lacking Spo11. Mol. Cell 6, 989–998.

Boekhout, M., Karasu, M.E., Wang, J., Acquaviva, L., Pratto, F., Brick, K., Eng, D.Y., Xu, J., Camerini-Otero, R.D., Patel, D.J., Keeney, S. (2019). REC114 Partner ANKRD31 Controls Number, Timing, and Location of Meiotic DNA Breaks. Mol. Cell 74, 1053–1068.

Boeri Erba, E., Signor, L., Oliva, M.F., Hans, F., Petosa, C. (2018). Characterizing intact macromolecular complexes using native mass spectrometry. Methods Mol. Biol. 1764, 133–151.

Boeri Erba, E., Signor, L., Petosa, C. (2020). Exploring the structure and dynamics of macromolecular complexes by native mass spectrometry. J. Proteomics. 222, 103799.

Brautigam, C.A. (2015). Calculations and Publication-Quality Illustrations for Analytical Ultracentrifugation Data. Methods in Enzymol. 562, 109–133.

Brinkmeier, J., Coelho, S., de Massy, B., Bourbon, H.-M. (2022). Evolution and Diversity of the TopoVI and TopoVI-like Subunits With Extensive Divergence of the TOPOVIBL subunit. Mol. Biol. Evol. 39, msac227.

Claeys Bouuaert, C., Pu, S., Wang, J., Oger, C., Daccache, D., Xie, W., Patel, D.J., Keeney, S. (2021a). DNA-driven condensation assembles the meiotic DNA break machinery. Nature 592. 144–149.

Claeys Bouuaert, C., Tischfield, S.E., Pu, S., Mimitou, E.P., Arias-Palomo, E., Berger, J.M., Keeney, S. (2021b). Structural and functional characterization of the Spo11 core complex. Nat. Struct. Mol. Biol. 28, 92–102.

de Massy, B. (2013). Initiation of Meiotic Recombination: How and Where? Conservation and Specificities Among Eukaryotes. Annu. Rev. Genet. 47, 563–599.

Dereli, I., Stanzione, M., Olmeda, F., Papanikos, F., Baumann, M., Demir, S., Carofiglio, F., Lange, J., De Massy, B., Baarends, W.M., Turner, J., Rulands, S., Tóth, A. (2021). Four-pronged negative feedback of DSB machinery in meiotic DNA-break control in mice. Nucleic Acids Res. 49, 2609–2628.

Henderson, K.A., Kee, K., Maleki, S., Santini, P.A., Keeney, S. (2006). Cyclin-Dependent Kinase Directly Regulates Initiation of Meiotic Recombination. Cell 125, 1321–1332.

Hunter, N. (2015). Meiotic recombination: The essence of heredity. Cold Spring Harb. Perspect. Biol. 7, a016618.

Jiao, K., Salem, L., Malone, R. (2003). Support for a Meiotic Recombination Initiation Complex: Interactions among Rec102p, Rec104p, and Spo11p. Mol. Cell. Biol. 23, 5928–5938.

Jumper, J., Evans, R., Pritzel, A., Green, T., Figurnov, M., Ronneberger, O., Tunyasuvunakool, K., Bates, R., Žídek, A., Potapenko, A., Bridgland, A., Meyer, C., Kohl, S.A.A., Ballard, A.J., Cowie, A., Romera-Paredes, B., Nikolov, S., Jain, R., Adler, J., Back, T., Petersen, S., Reiman, D., Clancy, E., Zielinski, M., Steinegger, M., Pacholska, M., Berghammer, T., Bodenstein, S., Silver, D., Vinyals, O., Senior, A.W., Kavukcuoglu, K., Kohli, P., Hassabis, D. (2021). Highly accurate protein structure prediction with AlphaFold. Nature 596, 583–589.

Kumar, R., Bourbon, H.M., De Massy, B. (2010). Functional conservation of Mei4 for meiotic DNA double-strand break formation from yeasts to mice. Genes Dev. 24. 1266–1280.

Kumar, R., Ghyselinck, N., Ishiguro, K. ichiro, Watanabe, Y., Kouznetsova, A., Höög, C., Strong, E., Schimenti, J., Daniel, K., Toth, A., de Massy, B. (2015). MEI4 - a central player in the regulation of meiotic DNA double-strand break formation in the mouse. J. Cell Sci. 128. 1800–1811.

Kumar, R., Oliver, C., Brun, C., Juarez-Martinez, A.B., Tarabay, Y., Kadlec, J., De Massy, B. (2018). Mouse REC114 is essential for meiotic DNA double-strand break formation and forms a complex with MEI4. Life Sci. Alliance. 1, e201800259.

Lam, I., Keeney, S. (2015). Mechanism and regulation of meiotic recombination initiation. Cold Spring Harb. Perspect. Biol. 7, a016634.

Maleki, S., Neale, M.J., Arora, C., Henderson, K.A., Keeney, S. (2007). Interactions between Mei4, Rec114, and other proteins required for meiotic DNA double-strand break formation in Saccharomyces cerevisiae. Chromosoma 116, 471–486.

Marty, M.T., Baldwin, A.J., Marklund, E.G., Hochberg, G.K.A., Benesch, J.L.P., Robinson, C. V. (2015). Bayesian deconvolution of mass and ion mobility spectra: From binary interactions to polydisperse ensembles. Anal. Chem. 87, 4370–4376.

Morgner, N., Robinson, C. V. (2012). Mass ign: An assignment strategy for maximizing information from the mass spectra of heterogeneous protein assemblies. Anal. Chem. 84, 2939–2948.

Nore, A., Juarez-Martinez, A.B., Clément, J., Brun, C., Diagouraga, B., Laroussi, H., Grey, C., Bourbon, H.M., Kadlec, J., Robert, T., de Massy, B. (2022). TOPOVIBL-REC114 interaction regulates meiotic DNA double-strand breaks. Nat. Commun. 13, 7048.

Panizza, S., Mendoza, M.A., Berlinger, M., Huang, L., Nicolas, A., Shirahige, K., Klein, F. (2011). Spo11-accessory proteins link double-strand break sites to the chromosome axis in early meiotic recombination. Cell 146, 372–383.

Papanikos, F., Clément, J.A.J., Testa, E., Ravindranathan, R., Grey, C., Dereli, I., Bondarieva, A., Valerio-Cabrera, S., Stanzione, M., Schleiffer, A., Jansa, P., Lustyk, D., Fei, J.F., Adams, I.R., Forejt, J., Barchi, M., de Massy, B., Toth, A. (2019). Mouse ANKRD31 Regulates Spatiotemporal Patterning of Meiotic Recombination Initiation and Ensures Recombination between X and Y Sex Chromosomes. Mol. Cell 74, 1069–1085.

Puglisi, R., Boeri Erba, E., Pastore, A. (2020). A Guide to Native Mass Spectrometry to determine complex interactomes of molecular machines. FEBS J. 287, 2428–2439.

Reinholdt, L.G., Schimenti, J.C. (2005). Mei1 is epistatic to Dmc1 during mouse meiosis. Chromosoma 114, 127–134.

Robert, T., Nore, A., Brun, C., Maffre, C., Crimi, B., Guichard, V., Bourbon, H.-M., de Massy, B. (2016). The TopoVIB-Like protein family is required for meiotic DNA double-strand break formation. Science. 351, 943–949.

Romanienko, P.J., Camerini-Otero, R.D. (2000). The mouse Spo11 gene is required for meiotic chromosome synapsis. Mol. Cell 6, 975–987.

Rousova, D., Nivsarkar, V., Altmannova, V., Raina, V.B., Funk, S.K., Liedtke, D., Janning, P., Müller, F., Reichle, H., Vader, G., Weir, J.R. (2021). Novel mechanistic insights into the role of Mer2 as the keystone of meiotic DNA break formation. Elife 10, e72330.

Sasanuma, H., Hirota, K., Fukuda, T., Kakusho, N., Kugou, K., Kawasaki, Y., Shibata, T., Masai, H., Ohta, K. (2008). Cdc7-dependent phosphorylation of Mer2 facilitates initiation of yeast meiotic recombination. Genes Dev. 22, 398–410.

Schuck, P. (2000). Size-distribution analysis of macromolecules by sedimentation velocity ultracentrifugation and Lamm equation modeling. Biophys. J. 78, 1606–1619.

Stanzione, M., Baumann, M., Papanikos, F., Dereli, I., Lange, J., Ramlal, A., Tränkner, D., Shibuya, H., de Massy, B., Watanabe, Y., Jasin, M., Keeney, S., Tóth, A. (2016). Meiotic DNA break formation requires the unsynapsed chromosome axis-binding protein IHO1 (CCDC36) in mice. Nat. Cell Biol. 18, 1208–1220.

Tessé, S., Bourbon, H.M., Debuchy, R., Budin, K., Dubois, E., Liangran, Z., Antoine, R., Piolot, T., Kleckner, N., Zickler, D., Espagne, E. (2017). Asy2/Mer2: An evolutionarily conserved mediator of meiotic recombination, pairing, and global chromosome compaction. Genes Dev. 31, 1880–1893.

Tock, A.J., Henderson, I.R. (2018). Hotspots for Initiation of Meiotic Recombination. Front. Genet. 9, 521.

Yadav, V.K., Claeys Bouuaert, C. (2021). Mechanism and Control of Meiotic DNA Double-Strand Break Formation in S. cerevisiae. Front. Cell Dev. Biol. 9, 642737.

