## Supplementary Information for "Characterization of the REC114-MEI4-IHO1 complex regulating meiotic DNA double-strand break formation"

Contains 13 Supplementary Figures

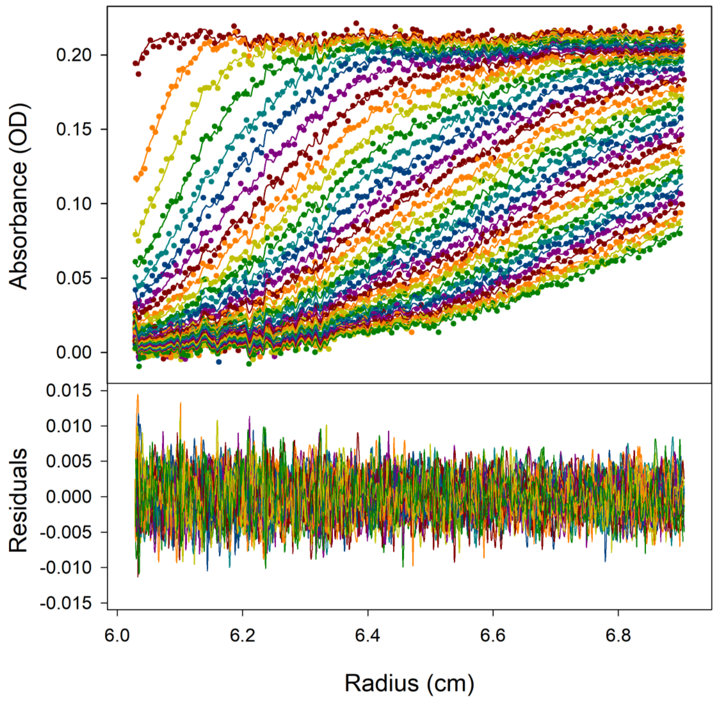


**Supplementary Figure 1**
Superimposition of experimental (dotted) and fitted (line) sedimentation velocity profiles, obtained at 280 nm, acquired every ~30 min during ~17h30 (top) and residuals (bottom), for sample at c = 0.2 mg/mL and pH 8.0.

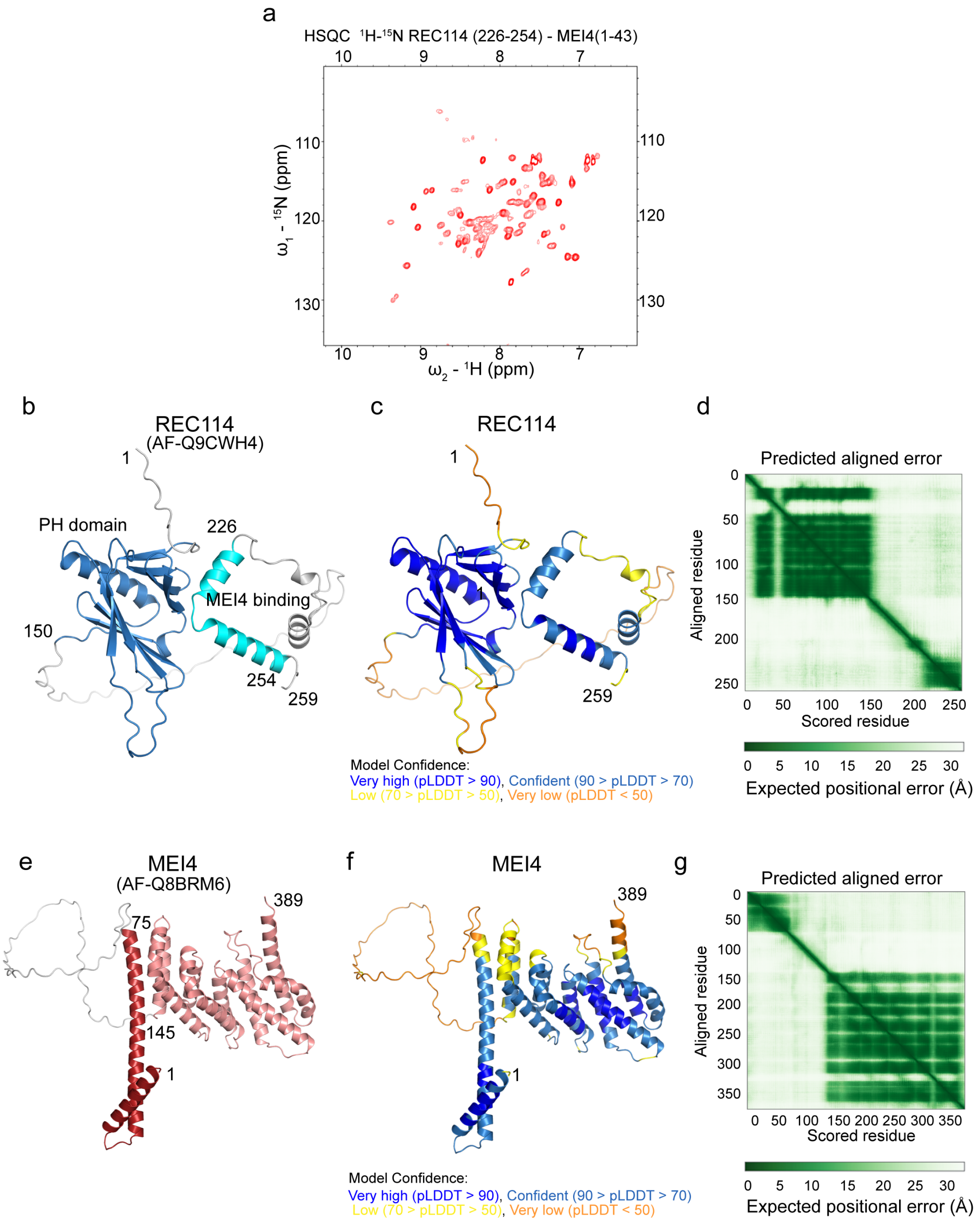


**Supplementary Figure 2**

**a.** $^{15}$N-HSQC spectrum of the REC114$^{226\text{-}254}$-MEI4$^{1\text{-}43}$ complex showing insufficient number of peaks that are not well resolved.

**b.** AlphaFold2 model of REC114 (AF-Q9CWH4) coloured by domain.

**c.** AlphaFold2 model of REC114 (AF-Q9CWH4) coloured according to the AlphaFold2 per-residue estimate of confidence (pLDDT).

**d.** Predicted aligned error plot for the model shown in **c**.

**e.** AlphaFold2 model of MEI4 (AF-Q8BRM6) coloured by domain.

**f.** AlphaFold2 model of REC114 (AF-Q8BRM6) coloured according to the AlphaFold2 per-residue estimate of confidence (pLDDT).

**g.** Predicted aligned error plot for the model shown in **e**.

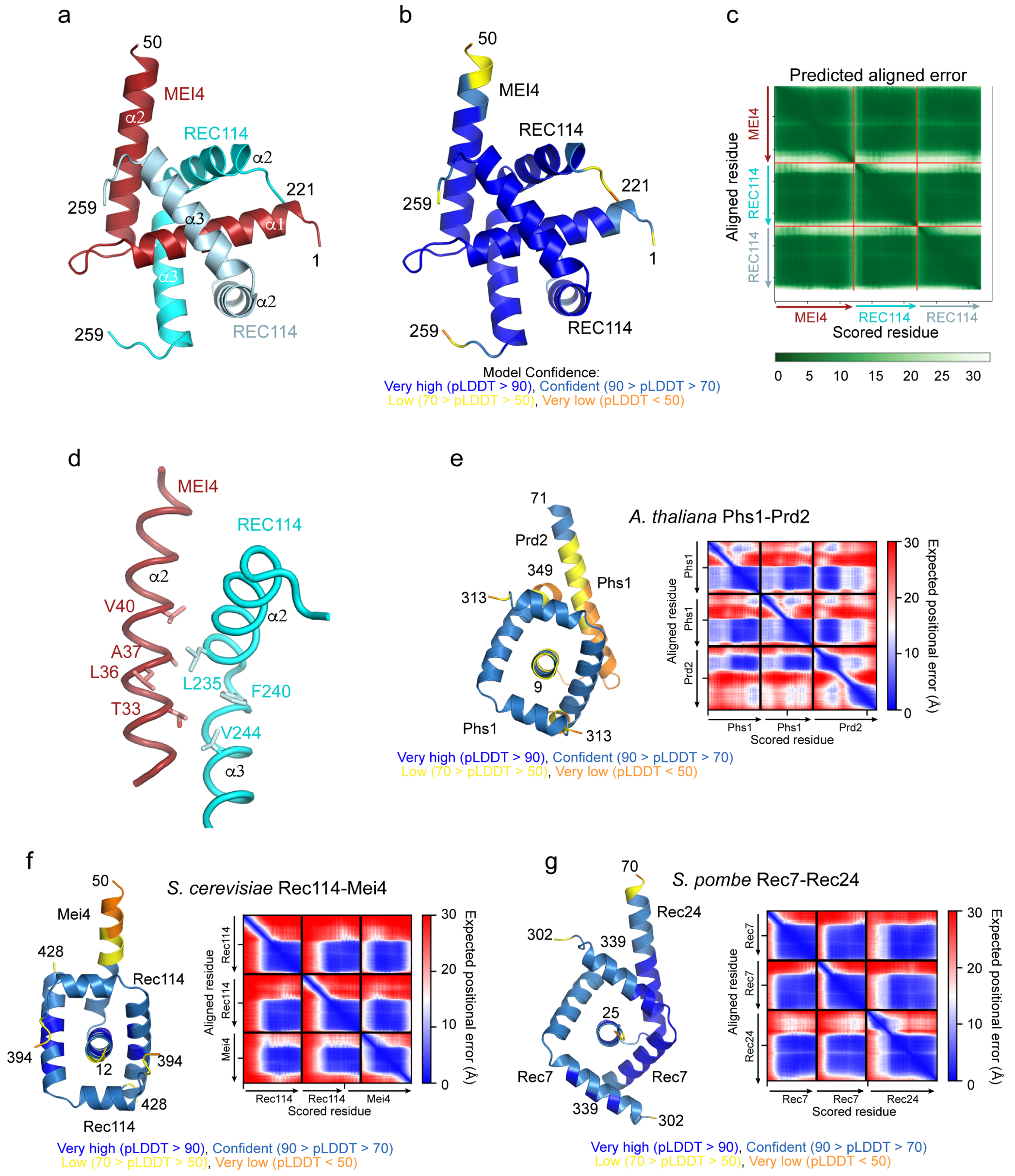


**Supplementary Figure 3**

**a.** AlphaFold2 model of the heterotrimeric complex between REC114$^{221\text{-}259}$-MEI4$^{1\text{-}50}$ coloured by protein chain.

**b.** AlphaFold2 model of the heterotrimeric complex between REC114$^{221\text{-}259}$-MEI4$^{1\text{-}50}$ coloured according to the AlphaFold2 per-residue estimate of confidence (pLDDT).

**c.** Predicted aligned error plot for the model shown in **a**.

**d.** Details of the predicted interaction between one of the REC114 molecules with the helix a2 of MEI4.

**e.** AlphaFold2 model coloured according to the AlphaFold2 per-residue estimate of confidence (pLDDT) and the predicted aligned error plot of the *A. thaliana* Phs1-Prd2 complex. The model was predicted for the first 80 residues of Prd2 and residues 284-349 of Phs1. Only residues 9-71 of Prd2 and 313-349 of Phs1 are shown as cartoon.

**f.** AlphaFold2 model coloured according to the AlphaFold2 per-residue estimate of confidence (pLDDT) and the predicted aligned error plot of *S. cerevisiae* Rec114-Mei4 complex. The model was predicted for the first 50 residues of Mei4 and residues 371-428 of Rec114. Only residues 12-50 of Mei4 and 394-428 of Rec114 are shown as cartoon.

**g.** AlphaFold2 model coloured according to the AlphaFold2 per-residue estimate of confidence (pLDDT) and the predicted aligned error plot of *S. pombe* Rec7-Rec24 complex. The model was predicted for the first 70 residues of Rec24 and residues 291-339 of Rec7. Only residues 25-70 of Rec24 and 302-339 of Rec7 are shown as cartoon.

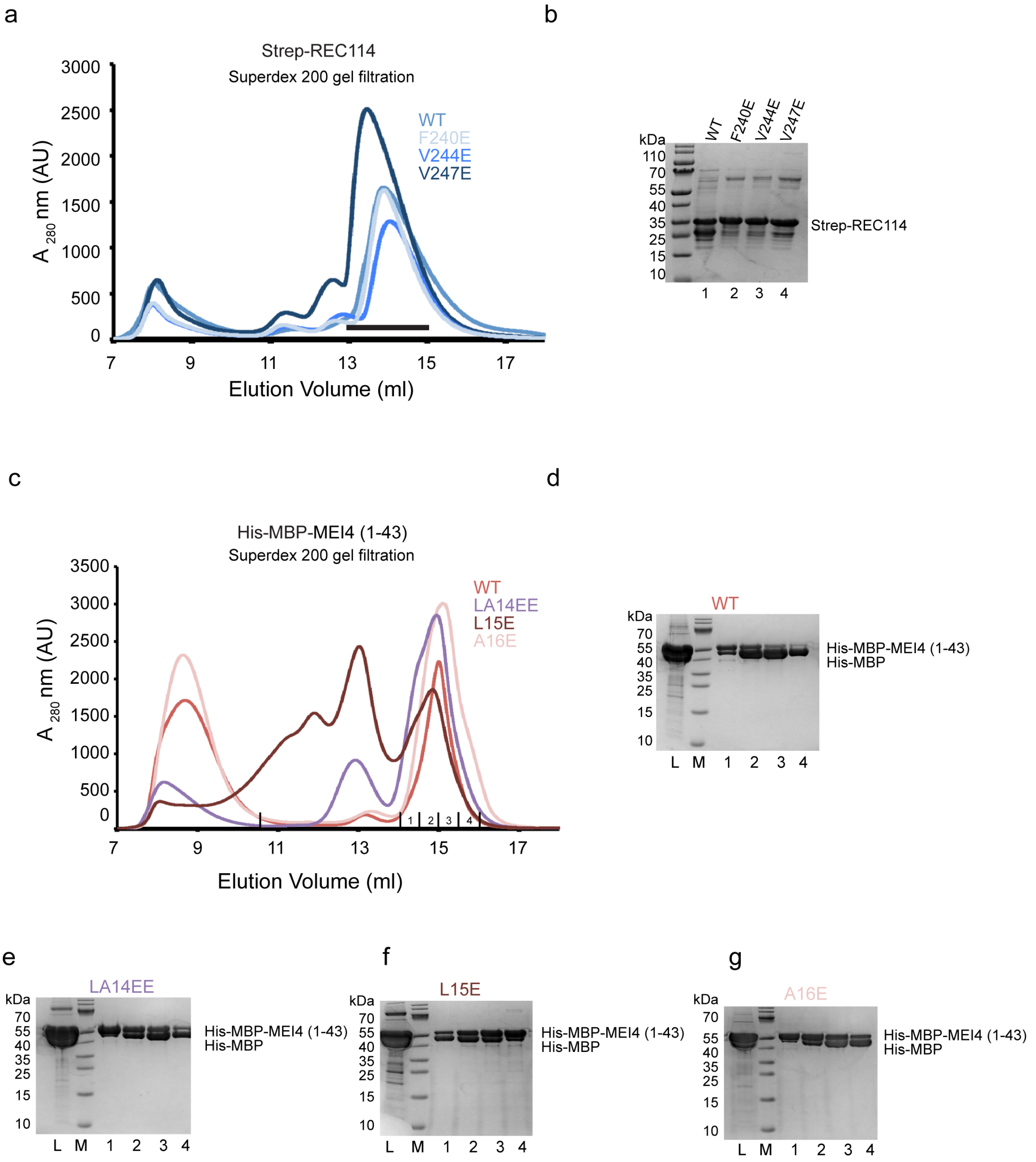


**Supplementary Figure 4**

**a.** Superdex 200 gel filtration elution profiles of WT and mutant Strep-REC114 proteins. The elution profiles are similar, indicating that the REC114 mutations do not significantly affect the overall structure of this REC114 fragment.

**b.** SDS-PAGE analysis of pooled fractions eluted between 13 and 15 ml of the Superdex 200 gel filtration elution profiles shown in **a**. Full-length REC114 is partially degraded, likely up to its PH domain.

**c.** Superdex 200 gel filtration elution profiles of WT and mutant His-MBP-MEI4$^{1-43}$ proteins. In absence of REC114, all the proteins have a tendency to aggregate.

**d-g.** SDS-PAGE analysis of fractions 1-4 of the Superdex 200 gel filtration elution profiles shown in **c**. L indicates input samples loaded onto the column. M indicates molecular weight marker.

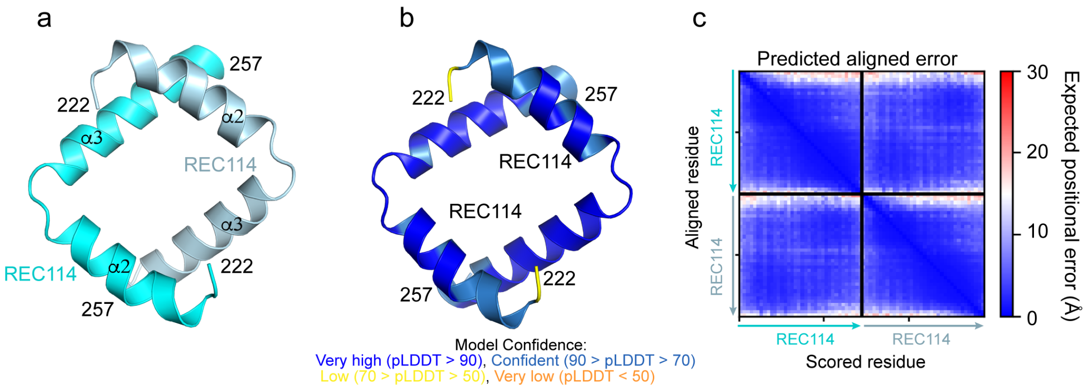


**Supplementary Figure 5**

**a.** AlphaFold2 model of the homodimerc complex of REC114$^{222\text{-}527}$ coloured by protein chain.

**b.** AlphaFold2 model of the homodimerc complex of REC114$^{222\text{-}257}$ coloured according to the AlphaFold2 per-residue estimate of confidence (pLDDT).

**c.** Predicted aligned error plot for the model shown in **a**.

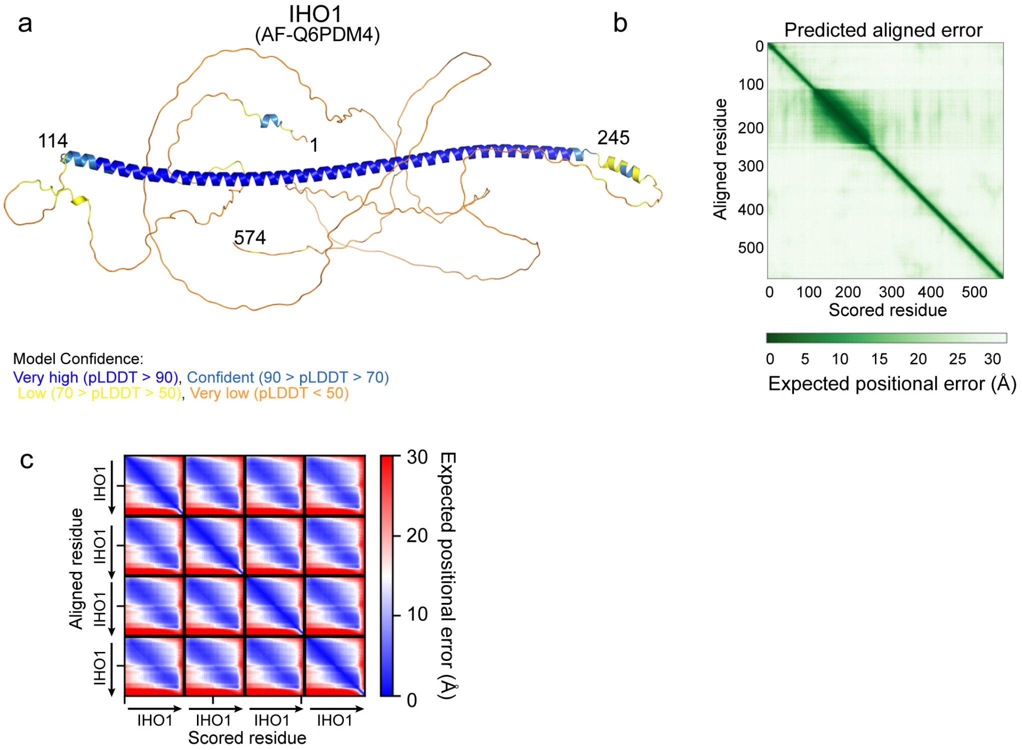


**Supplementary Figure 6**

**a.** AlphaFold2 model of IHO1 (AF-Q6PDM4) coloured according to the AlphaFold2 per-residue estimate of confidence (pLDDT).

**b.** Predicted aligned error plot for the model shown in **c**.

**c.** Predicted aligned error plot for the model shown in Fig. 4c.

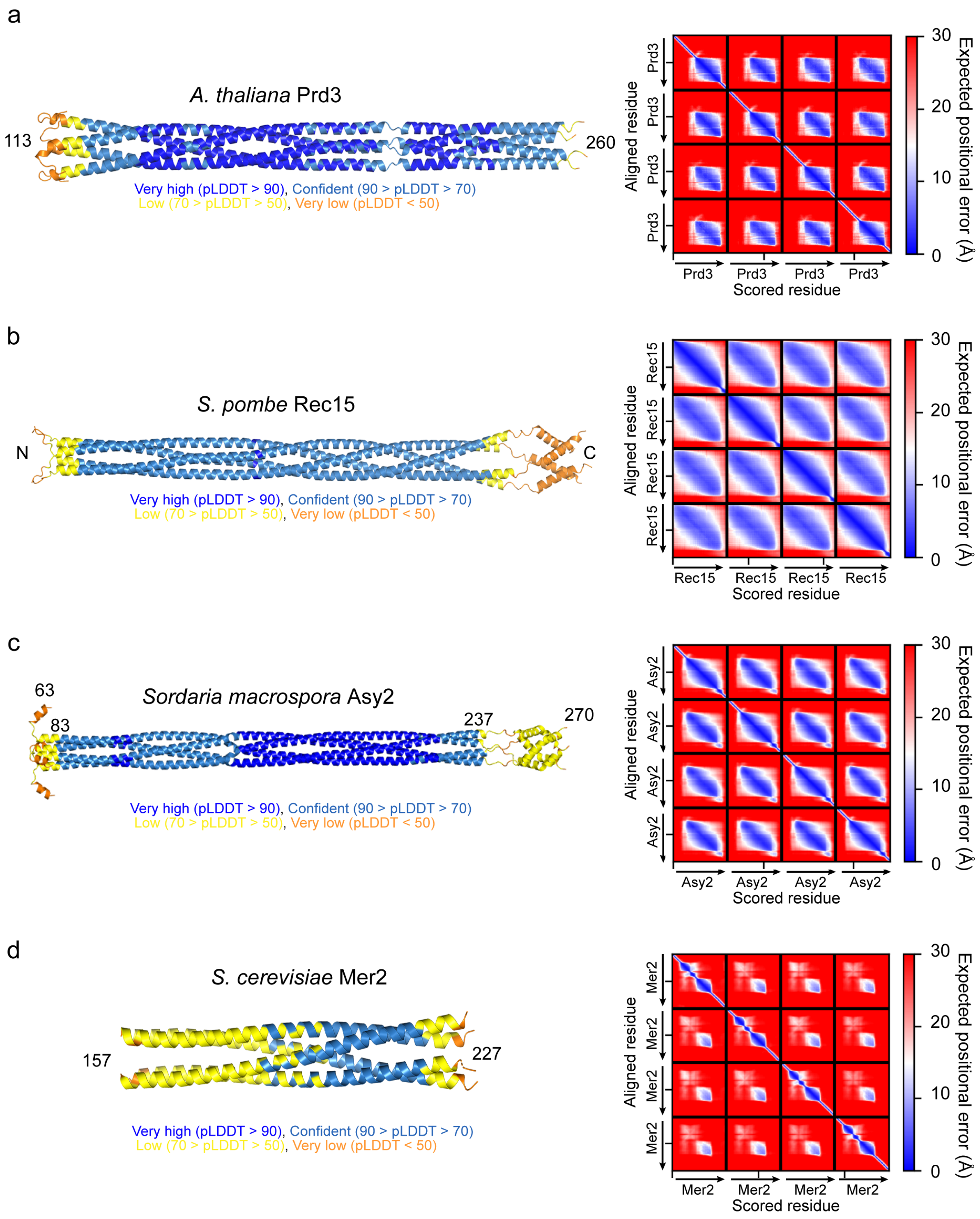


**Supplementary Figure 7**

**a.** AlphaFold2 model coloured according to the AlphaFold2 per-residue estimate of confidence (pLDDT) and the predicted aligned error plot of *A. thaliana* Prd3. The model was predicted for the first 300 residues and only residues 113-260 are shown as cartoon.

**b.** AlphaFold2 model coloured according to the AlphaFold2 per-residue estimate of confidence (pLDDT) and the predicted aligned error plot of *S. pombe* Rec15.

**c.** AlphaFold2 model coloured according to the AlphaFold2 per-residue estimate of confidence (pLDDT) and the predicted aligned error plot of *Sordaria mocrospora* Asy2. The model was predicted for the first 300 residues and only residues 63-270 are shown as cartoon.
**d.** AlphaFold2 model coloured according to the AlphaFold2 per-residue estimate of confidence (pLDDT) and the predicted aligned error plot of *S.cerevisiae* Mer2. The model was predicted for the full-length protein and only residues 157-227 are shown as cartoon.

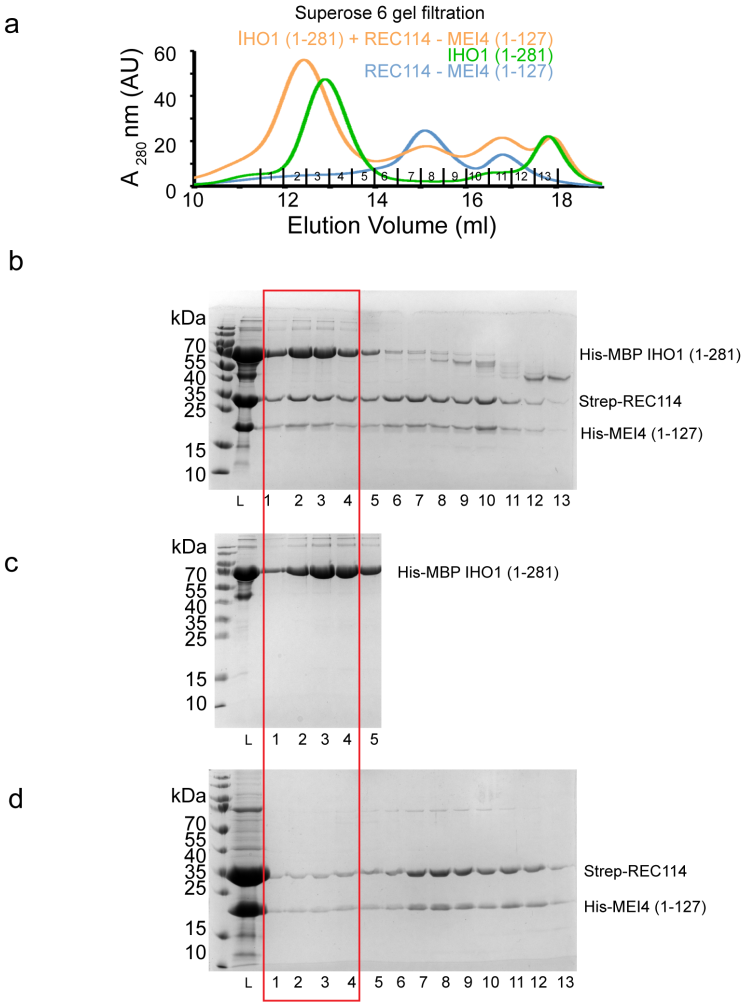


**Supplementary Figure 8**

**a.** Overlay of Superose 6 gel filtration elution profiles of His-MBP-IHO1$^{1-281}$, the Strep-REC114-His-MEI4$^{1-127}$ complex and their mixture.

**b-d**. SDS-PAGE analysis of fraction 1-13 of the Superose 6 gel filtration elution profile shown in **a**. Only fractions 1-5 are shown for the His-MBP-IHO1$^{1-281}$ profile. L indicates input sample loaded onto the column.

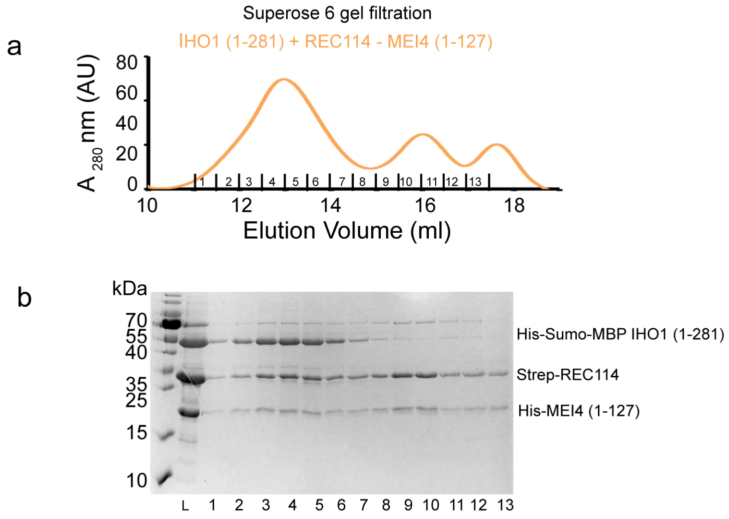


**Supplementary Figure 9**

**a.** Superose 6 gel filtration elution profile of His-Sumo-IHO1$^{1\text{-}281}$ mixed with the Strep-REC114-His-MEI4$^{1\text{-}127}$ complex.

**b**. SDS-PAGE analysis of fraction 1-13 of the Superose 6 gel filtration elution profile shown in **a**. L indicates input sample loaded onto the column.

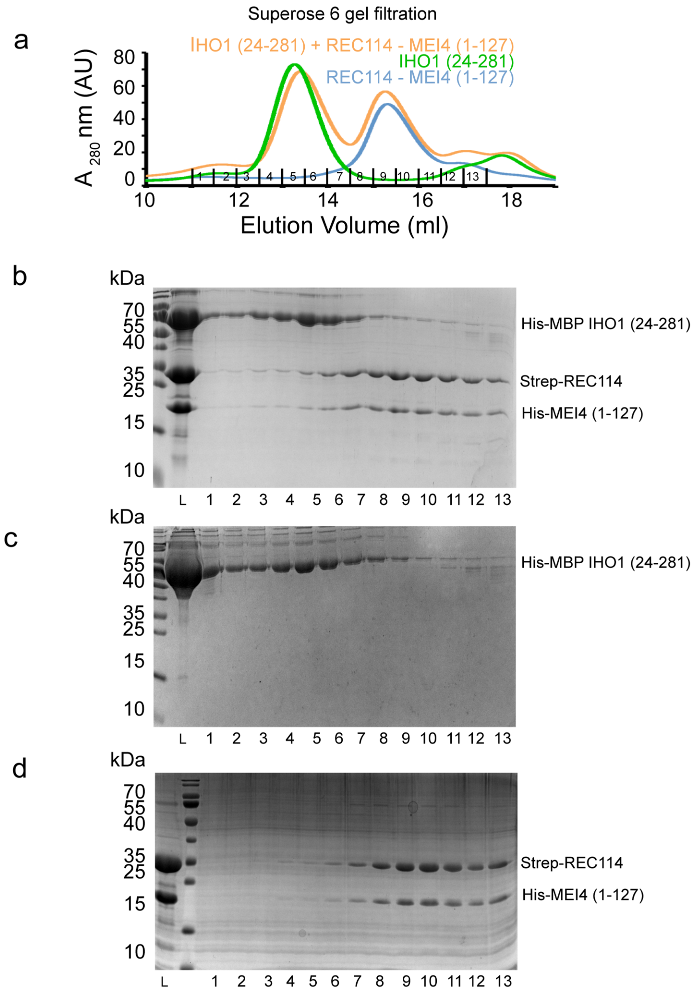


**Supplementary Figure 10**

**a.** Overlay of Superose 6 gel filtration elution profiles of His-MBP-IHO1$^{24\text{-}281}$, the Strep-REC114-His-MEI4$^{1\text{-}127}$ complex and their mixture.

**b-d**. SDS-PAGE analysis of fraction 1-13 of the Superose 6 gel filtration elution profile shown in **a**. L indicates input sample loaded onto the column.

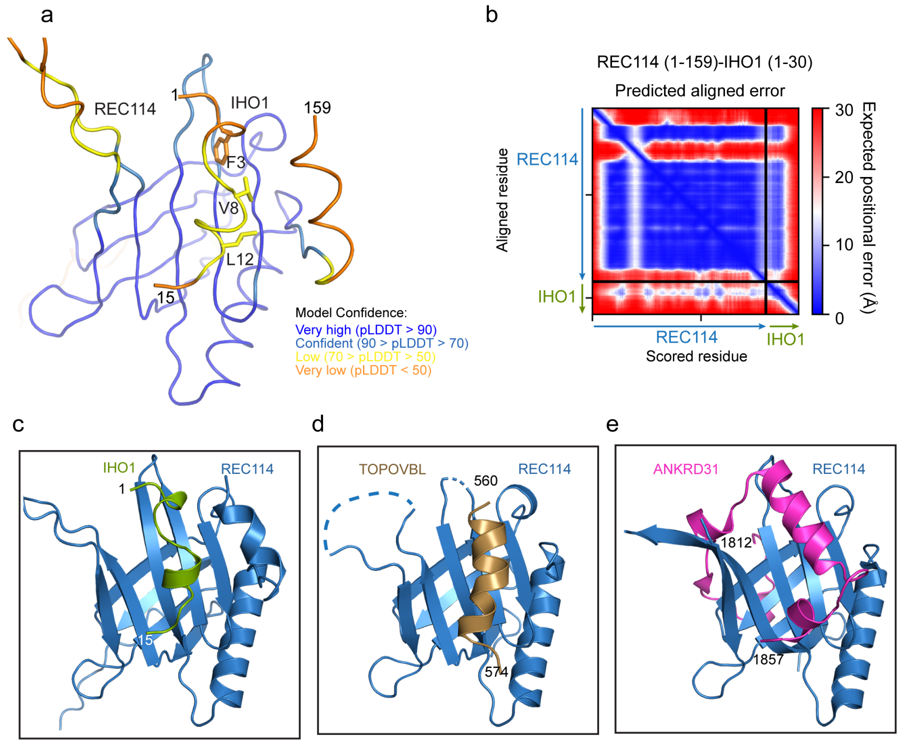


**Supplementary Figure 11**

**a.** AlphaFold2 model of the complex between REC114$^{1-159}$ and IHO1$^{1-30}$ coloured according to the AlphaFold2 per-residue estimate of confidence (pLDDT). Only IHO1 residues 1-15 are shown.

**b.** Predicted aligned error plot for the model shown in **a**.

**c-e.** Comparison of the AlphaFold2 model of the complex between the PH domain of REC114 and IHO1$^{1-15}$ with crystal structure of the REC114 PH domain bound to a peptide of TOPOVIBL (PDB code: 7QWV) and ANKRD31 (PDB code: 6NFX).

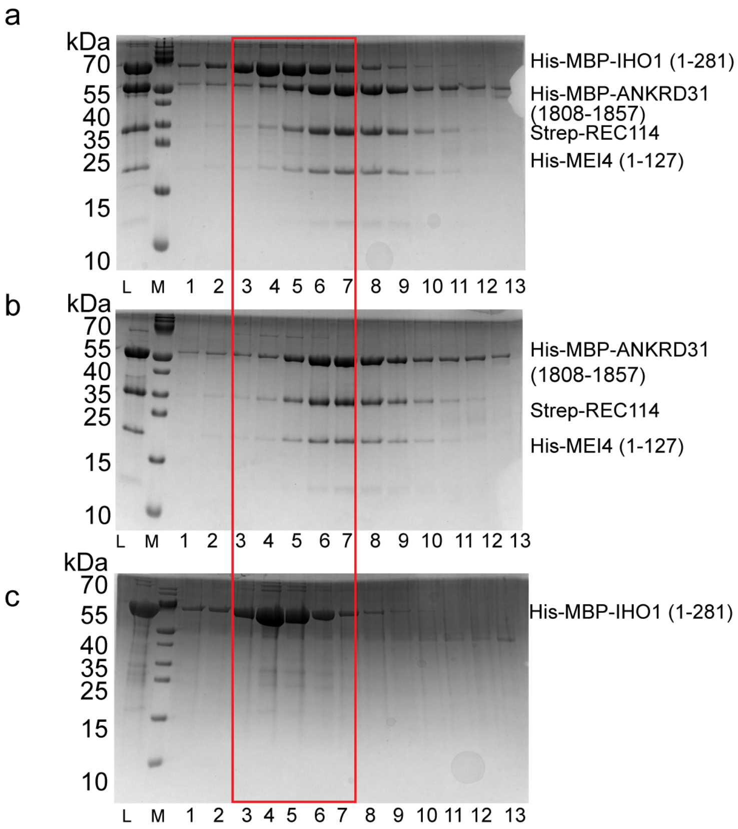


**Supplementary Figure 12**

**a-c.** SDS-PAGE analysis of fractions 1-13 of Superose 6 gel filtration elution profiles shown in Fig. 6i.
**a.** Elution profile of His-MBP-IHO1$^{1\text{-}281}$ mixed with His-MBP-ANKRD31$^{1808\text{-}1857}$, Strep-REC114 and His-MEI4$^{1\text{-}127}$. L indicates input sample loaded onto the column.
**b.** Elution profile of His-MBP-ANKRD31$^{1808\text{-}1857}$ bound to Strep-REC114-His-MEI4$^{1\text{-}127}$.
**c.** Elution profile of His-MBP-IHO1$^{1\text{-}281}$.
The red frame highlights essentially the same position of His-MBP-IHO1$^{1\text{-}281}$ when eluted alone or when mixed with His-MBP-ANKRD31$^{1808\text{-}1857}$ and Strep-REC114-His-MEI4$^{1\text{-}127}$.

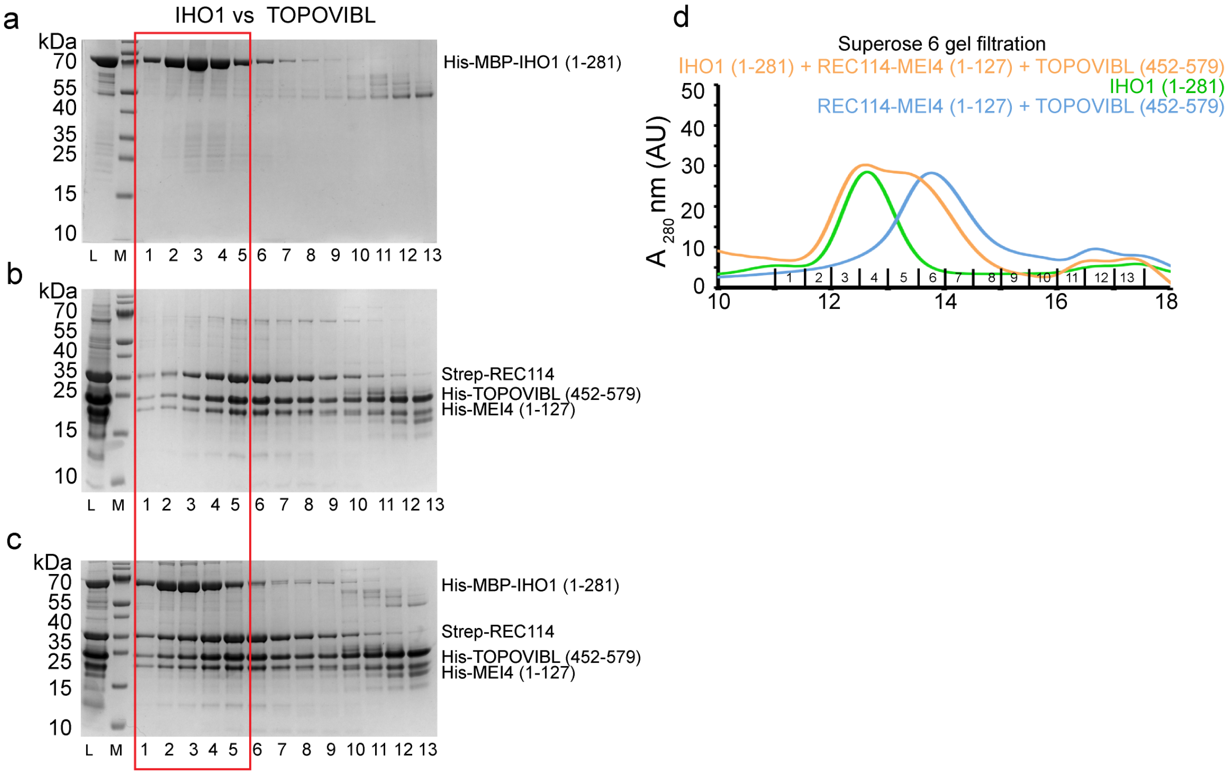


**Supplementary Figure 13**

**a-c.** SDS-PAGE analysis of fractions 1-13 of Superose 6 gel filtration elution profiles shown in **d**.

**a.** Elution profile of His-MBP-IHO1$^{1-281}$. L indicates input sample loaded onto the column.

**b.** Elution profile of His-TOPOVIBL$^{452-579}$ bound to Strep-REC114-His-MEI4$^{1-127}$ .

**c.** Elution profile of His-MBP-IHO1$^{1-281}$ mixed with His-TOPOVIBL$^{452-579}$, Strep-REC114 and His-MEI4$^{1-127}$. The red frame highlights essentially the same position of His-MBP-IHO1$^{1-281}$ when eluted alone or when mixed with His-TOPOVIBL$^{452-579}$ and Strep-REC114-His-MEI4$^{1-127}$.

**d.** Superose 6 gel filtration elution profiles of His-MBP-IHO1$^{1-281}$ mixed with His-TOPOVIBL$^{452-579}$, Strep-REC114 and His-MEI4$^{1-127}$ (orange); His-TOPOVIBL$^{452-579}$, Strep-REC114 and His-MEI4$^{1-127}$ (blue) and His-MBP-IHO1$^{1-281}$ (green).
